# Metabolic Reprogramming Driven by Ant2 -Targeting Augments T-Cell Function and Anti-Tumor Immunity

**DOI:** 10.1101/2024.04.21.590436

**Authors:** Omri Yosef, Leonor Cohen-Daniel, Oded Shamriz, Zahala Bar-On, Amijai Saragovi, Ifat Abramovich, Bella Agranovich, Veronika Lutz, Joseph Tam, Anna Permyakova, Eyal Gottlieb, Magdalena Huber, Michael Berger

**Author notes:** Correspondence (M.B.).

## Abstract

T-cell activation requires a substantial increase in NAD^+^ production, often exceeding the capacity of oxidative phosphorylation (OXPHOS). To investigate how T cells adapt to this metabolic challenge, we generated T cell-specific ADP/ATP translocase-2 knockout (Ant2^−/−^) mice. Loss of Ant2, a crucial protein mediating ADP/ATP exchange between mitochondria and cytoplasm, induces OXPHOS restriction by limiting ATP synthase activity, impeding NAD^+^ regeneration. Interestingly, Ant2^−/−^ naïve T cells exhibited enhanced activation, proliferation, and effector functions compared to wild-type controls. Metabolic profiling revealed that these cells adopt an activated-like metabolic program with increased mitobiogenesis and anabolism. Pharmacological inhibition of ANT in wild-type T cells recapitulated the Ant2^−/−^ phenotype, and improved adoptive cell therapy of cancer. Our findings suggest that Ant2-deficient T cells bypass the typical metabolic reprogramming required for activation, leading to enhanced T-cell function. These results highlight the critical role of mitochondrial metabolism in regulating T-cell fate and underscore the therapeutic potential of targeting ANT for immune modulation.

## Introduction

Metabolic rewiring in T cells plays a pivotal role in regulating their activation, fate decision, and overall immune function.^1–5^ Upon activation, T cells undergo substantial metabolic reprogramming to meet the increased demands for raw energy (ATP), potential energy (NAD^+^), and biosynthetic precursors.^1^ This metabolic rewiring involves the upregulation of aerobic glycolysis and oxidative phosphorylation (OXPHOS).^6^ As the metabolic hub of the cell,^7^ the mitochondrion undergoes extensive remodeling during activation to support anabolism, including increased biomass, morphological changes, augmented ROS production, and altered metabolic pathways within minutes of T cell receptor (TCR) ligation.^8,9^

Rapidly proliferating cells, such as activated T cells, exhibit an elevated demand for NAD^+^. This increased NAD^+^ requirement presents a significant metabolic bottleneck as NAD^+^ regeneration is intrinsically linked to ATP production via OXPHOS. The rate-limiting step in this process is the activity of ATP synthase, which dictates the pace of NAD^+^ regeneration. Consequently, OXPHOS’ capacity to meet the heightened NAD^+^ demands of these cells is constrained, resulting in OXPHOS insufficiency.^10^

Our previous findings indicate that in effector T cells, OXPHOS primarily supports anabolic tricarboxylic acid (TCA) cycle by NADH oxidation, rather than ATP production, emphasizing the critical role of NAD^+^ regeneration.^11^ This is supported by the observation that T cells can function under hypoxic conditions,^12,13^ indicating that mitochondrial ATP (mtATP) generation is dispensable for T-cell functions. To understand how T cells adapt to this metabolic bottleneck, we focused on naïve T cells. We hypothesized that these cells might employ specific strategies to maintain cellular homeostasis under conditions of constrained OXPHOS, which may influence the transition to an activated state.

To evaluate this possibility, we indirectly induced chronic OXPHOS restriction by generating T cell-specific ADP/ATP translocase-2 knockout (Ant2^−/−^) mice. Ant2, encoded by the *Slc25a5* gene, transports ADP/ATP between the mitochondria and the cytosol, maintaining mitochondrial membrane potential (ΔΨ), and supporting cellular energy homeostasis.^14,15^ Ant2 deficiency reduces mtATP transfer to the cytosol, accompanied by a concomitant decrease in ADP concentrations within the mitochondrial matrix. Consequently, ATP-synthase is unable to dissipate the ΔΨ due to reduced ADP availability. This sustained perturbation negatively affects the efficiency of the electron transport chain (ETC), thereby restricting OXPHOS and NAD^+^ regeneration, effectively mimicking the OXPHOS insufficiency experienced by activated T cells. By studying these cells, we aimed to determine whether and how these adaptations influence subsequent activation and effector functions.

Here, we observed that T-cell-specific Ant2^−/−^ mice experienced T-cell lymphopenia due to an inability to respond to homeostatic expansion signals. Surprisingly, Ant2-deficient T cells exhibited enhanced activation propensity, proliferation and effector functions compared to wild-type (WT) controls. Proteomic and metabolic analyses revealed an activated-like metabolic signature with increased anabolism and distinct mitochondrial characteristics. In line with our results, pharmacological inhibition of ANT activity enhanced naïve WT T cell responsiveness to activation stimuli, with increased mitobiogenesis and improved proliferation and anti-tumor activity in different *in-vivo* models. We propose that naïve Ant2^−/−^ T cells bypass the typical metabolic reprogramming during priming, resulting in activated-like traits and behavior. Our study identifies Ant2 as a critical regulator of T cell metabolic plasticity and provides a mechanistic basis for developing therapeutic interventions targeting mitochondrial metabolism to enhance immune responses.

## Results

### Ant2^−/−^ leads to T-cell lymphopenia due to failure to respond to homeostatic expansion signals

To test the impact of depleting mitochondrial ATP from T cells, we generated T-cell-specific Ant2^−/−^ mice by crossbreeding Ant2^fl/fl^ mice with distal Lck-cre transgenic mice (dLck-Cre^+^). The resulting mice, referred to as Ant2^−/−^, had *Slc25a5* E2+3 deleted in CD3^+^ cells, confirmed through polymerase chain reaction (PCR) (Figure S1A-B). This deletion resulted in null alleles, as Ant2 mRNA was barely detectable in reverse-transcriptase PCR (RT-PCR) (Figure S1C). Notably, Ant1 mRNA levels remained similar between Ant2^−/−^ and WT cells. Immunoblot analysis using an anti-Ant2 antibody further confirmed the absence of Ant2 protein expression in Ant2^−/−^ T cells (Figure S1D).

To investigate the immunological effects of Ant2 deficiency, we conducted immunophenotyping using flow cytometry. Ant2^−/−^ mice exhibited normal cellularity in the spleen, thymus, lymph nodes, and peripheral blood (data not shown). Reduced percentages and counts of CD4^+^ and CD8^+^ T cells were evident in both the spleen (Figure 1A-B) and lymph nodes (Figure S1E-F), indicating that Ant2^−/−^ mice exhibit T-cell lymphopenia. In addition, Ant2^−/−^ CD8^+^ T cells showed an increase in the memory-like subpopulations, characterized by elevated levels of CD44^+^CD62L^+^ (Figure S1G-H) and CD44^+^CD122^+^ (Figure S1I-J) subpopulations, at the expense of naïve subpopulations (CD62L^+^CD44^−^ and CD122^−^CD44^−^). Notably, Ant2^−/−^ CD8^+^ T cells displayed intact CD127 (IL7-R) (Figure S1K-L). These results strongly suggest that the Ant2^−/−^ T cells acquire a memory-like phenotype due to lymphopenia-induced homeostasis-stimulated proliferation.^16,17^

**Figure 1:**
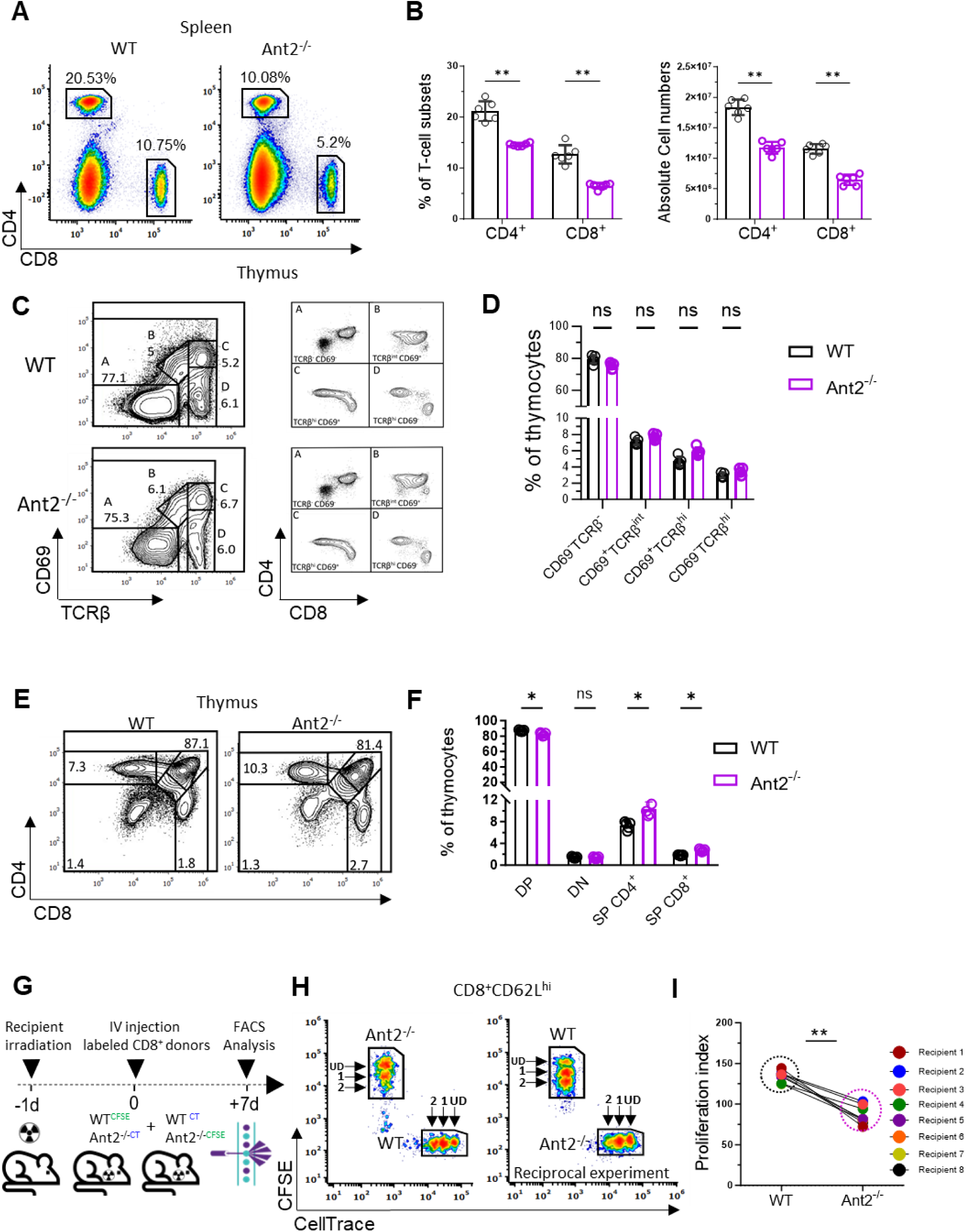
Impaired homeostatic expansion of Ant2^−/−^ T cells. **(A)** Representative flow cytometry plots showing the frequency of CD4+ and CD8^+^ T cells in the spleen from WT and Ant2^−/−^ littermate mice (6-9 weeks old). **(B)** Bar graphs summarizing the results in A, indicating percentages **(left)** and absolute cell numbers **(right)**. **(C-F)** Analysis of thymus development of WT and Ant2^−/−^ littermate mice (6-9 weeks old). **(C) Left panel:** representative flow cytometry plots showing four developmental stages (A-D), distinguished by CD69 vs TCRβ staining of whole thymocytes. **Right panel:** CD4 vs. CD8 staining gated on each of the developmental stages, A-D. **(D)** Bar graph summarizing the results shown in C left panel. **(E)** Representative flow cytometry plots showing the frequency of CD4 vs. CD8 in the thymus-derived from 6-9 weeks old WT and Ant2^−/−^ littermate mice. **(F)** Bar graph summarizing the results of panel F. DP-double positive, DN-double negative, and SP-single positive. **(G)** Schematic of homeostatic expansion experiment; co-adoptive transfer of CFSE-labeled WT, and CellTrace-labeled Ant2^−/−^ -derived splenocytes to sub-lethally irradiated recipient mice. The reciprocal experiment was also performed. **(H)** Representative flow cytometry plots showing CFSE vs CellTrace intensities gated on donor CD8^+^ CD62L^hi^ T cells. **(I)** Graph depicting paired proliferation indexes of donor WT and Ant2^−/−^ CD8^+^ T cells. Statistical method: Two-tailed Mann–Whitney test (B, D, F), or two-tailed Wilcoxon matched-pairs signed rank test (I). Data are represented as mean ± S.D. (P value, *P≤0.05 **P≤0.01).

To understand the underlying causes of T-cell lymphopenia in Ant2^−/−^ mice, we initially examined thymic development using flow cytometry. Comparing WT and Ant2^−/−^ mice, we found that the pre- and post-positive selection thymic populations, CD69^−^TCRβ^−^ and CD69^+^TCRβ^hi^, respectively, were comparable between the two groups (Figure 1C-D).^18^ However, we observed a slight but significant decrease in the double-positive (CD4^+^CD8^+^) cell population and an increase in the single-positive (CD4^+^ or CD8^+^) cell populations in Ant2^−/−^ mice (Figure 1E-F). These findings suggest intact thymic development with a minor increase in thymic egress, which is consistent with the observed T-cell lymphopenia in Ant2^−/−^ mice.

To evaluate the expansion capacity of naïve Ant2^−/−^ T cells in the secondary lymphatic system, we conducted a proliferation assay in response to homeostatic expansion signals. We performed adoptive transfer experiments using CD45.2^+^ CellTrace-labeled WT splenocytes and carboxyfluorescein diacetate succinimidyl ester (CFSE)-labeled Ant2^−/−^ splenocytes. The transferred cells were introduced into sub-lethally irradiated syngeneic recipient mice (CD45.1^+^). Additionally, a reciprocal experiment was conducted using CFSE-labeled WT splenocytes and CellTrace-labeled Ant2^−/−^ splenocytes. Flow cytometry analysis of donor T cells was performed 7 days post-transfer (Figure 1G). Our results demonstrate that Ant2^−/−^ CD8^+^ (Figure 1H-I) and CD4^+^ (Sup 1M-N) T cells exhibited significantly reduced proliferation compared to WT cells. Collectively, these findings strongly suggest that the T-cell lymphopenia observed in Ant2^−/−^ mice is primarily attributed to a defect in the expansion of naïve T cells in the secondary lymphatic system, rather than impaired thymic development.

### Ant2^−/−^ mice demonstrate intact T-cell-dependent immunity

To assess the impact of T-cell lymphopenia on T-cell-dependent immunity in Ant2^−/−^ mice, we performed several *in-vivo* experiments. First, we assessed CD4^+^ T-cell immunity by examining the thymus-dependent B cell antibody response and cytokine production in mice immunized with ovalbumin (OVA) (Figure 2A).^19^ Surprisingly, the concentrations of OVA-specific IgG1 and IgG2a antibodies (Figure 2B-C), as well as the cytokines IFN-γ and IL-4 (Figure 2D-E), were found to be equivalent in the sera of both WT and Ant2^−/−^ immunized mice.

**Figure 2:**
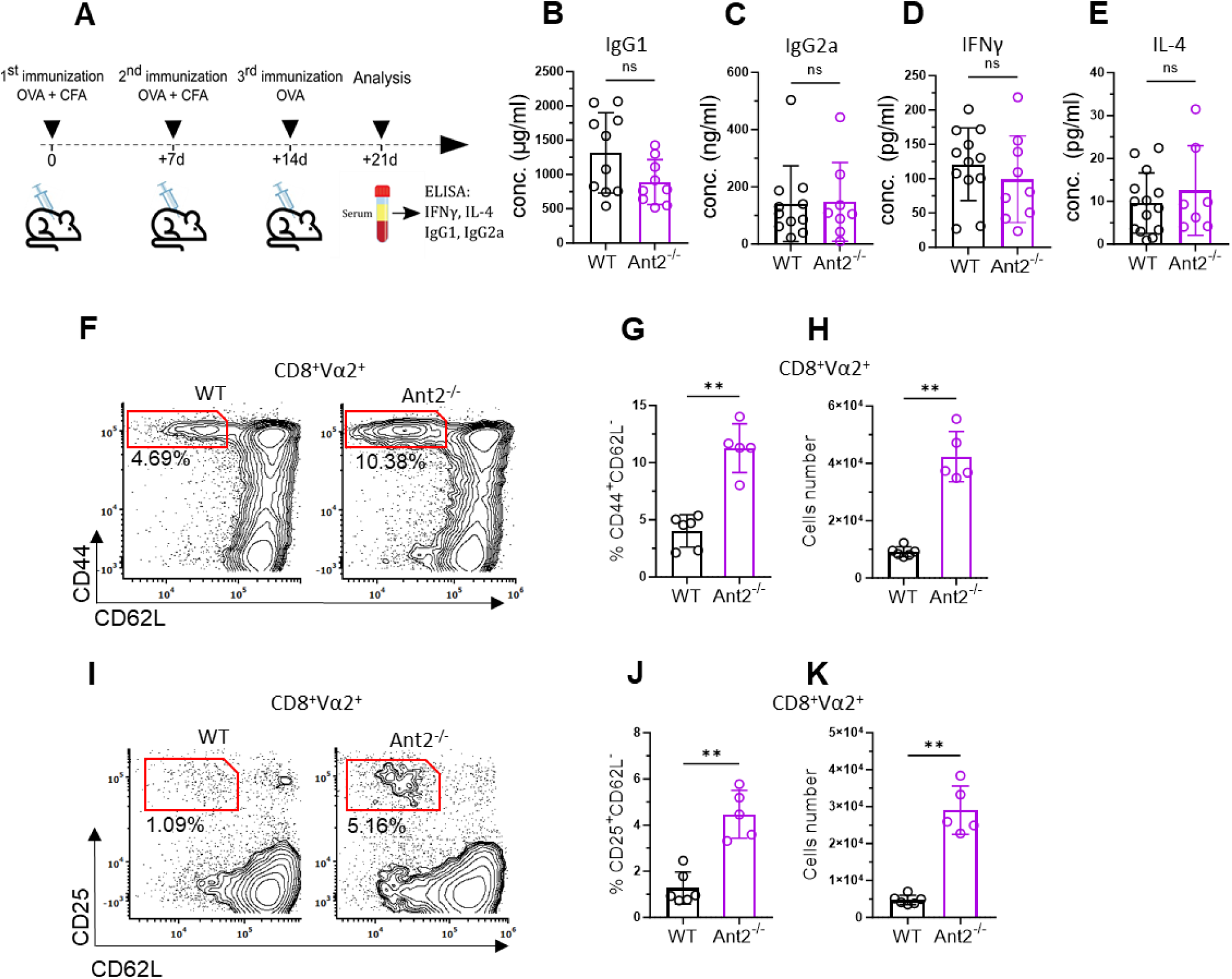
Intact T-cell mediated immunity in Ant2^−/−^ mice. **(A)** Schematic representation of the immunization experiment used to estimate CD4^+^ T-cell responses. WT and Ant2^−/−^ mice were immunized three times at 7-day intervals. The first two immunizations included OVA protein in complete Freund’s adjuvant (CFA), while the third included OVA without CFA. One week after the third injection the concentration of, OVA-specific IgG1 **(B)** and IgG2a **(C)**, as well as the cytokines IFNγ **(D)** and IL-4 **(E)**, were measured in the serum of the mice using enzyme-linked immunosorbent assay (ELISA). **(F-K)** WT and Ant2^−/−^ littermate mice were primed intradermally in the left ear-pinna with 5 × 10^6^ transduction units (TU) of lentivirus expressing OVA (LvOVA). Seven days post-infection, cervical lymph nodes (LN) were dissected, and single cells were stained for activation-related surface markers. **(F)** Representative flow cytometry plots showing CD62L vs. CD44 gated on CD8^+^Vα2^+^ T cells. **(G)** Bar graph summarizing the results shown in F. **(H)** Absolute cell number of CD8^+^Vα2^+^CD44^+^CD62L^−^ in the cervical LN of WT and Ant2^−/−^ mice. **(I)** Representative flow cytometry plots showing CD62L vs. CD25 gated on CD8^+^Vα2^+^ T cells. **(J):** Bar graph summarizing the results shown in I. **(K)** Absolute cell number of CD8^+^Vα2^+^CD25^+^CD62L^−^ in the cervical LN of WT and Ant2^−/−^ mice. Statistical method: Two-tailed Mann–Whitney test. Data are represented as mean ± S.D. (P value, **P≤0.01).

Next, to evaluate CD8^+^ T cell function, we intradermally injected WT and Ant2^−/−^ mice with an OVA-expressing lentivirus (Lv-OVA) into the left ear pinna.^11,20^ After seven days, we assessed the activation status and ratios of OVA-associated CD8^+^ T cells (TCR Vα2^+^) in the left deep cervical lymph nodes (LN), using the right LN as a control. Activation status was determined by staining for established *in-vivo* activation markers, including elevation of CD44 and CD25, and reduction of CD62L. Notably, the left LN of all tested mice exhibited visible swelling and an increase in absolute cell number compared to their right LN (Figure S2A-B). Consistent with the observed lymphopenia, Ant2^−/−^ left LN demonstrated a decrease in the total CD8^+^ ratios compared to WT (Figure S2C-D). However, the percentages of OVA-associated cells (CD8^+^Va2^+^) were comparable between the two groups (Figure S2C, E). Furthermore, CD8^+^Vα2^+^ cells derived from Ant2^−/−^ mice showed an increase in both the percentages and absolute numbers of CD62L^−^CD44^+^ (Figure 2F-H) and CD62L^−^CD25^+^ (Figure 2I-K) cells compared to their WT counterparts.

In summary, our findings indicate that the observed lymphopenia in Ant2^−/−^ mice does not significantly impact their T cell-dependent immune responses. Instead, we observed an increase in effector markers in CD8^+^ T cells. This increase in activation markers may explain the intact concentrations of cytokines in the sera of Ant2^−/−^ mice, suggesting T-cell hyperresponsiveness as a compensatory mechanism to preserve T-cell-dependent immunity.

### Ant2^−/−^ T cells have a higher propensity to activation stimuli and improved effector functions

Based on our *in-vivo* results, we theorized that Ant2^−/−^ T cells might demonstrate increased sensitivity to activation cues and heightened effector functions. To test our hypothesis, we conducted several *in-vitro* experiments. We examined their response to different concentrations of αCD3/αCD28 antibodies, which allowed us to assess their activation, proliferation capacities, and effector function. Activated Ant2^−/−^ CD8^+^ T cells exhibited enhanced proliferation capacity compared to their WT counterpart across all tested concentrations of αCD3/αCD28 antibodies (Figure 3A-B). Notably, expression of CD25 was comparable between the groups (Figure 3C-D). Moreover, we observed a significant increase in both the proportions of CD25^+^IFNγ^+^ cells among total CD8^+^ cells and the intensity of IFNγ per cell in Ant2^−/−^ CD8^+^ T cells when compared to their WT controls (Figure 3C, 3E-F).

**Figure 3:**
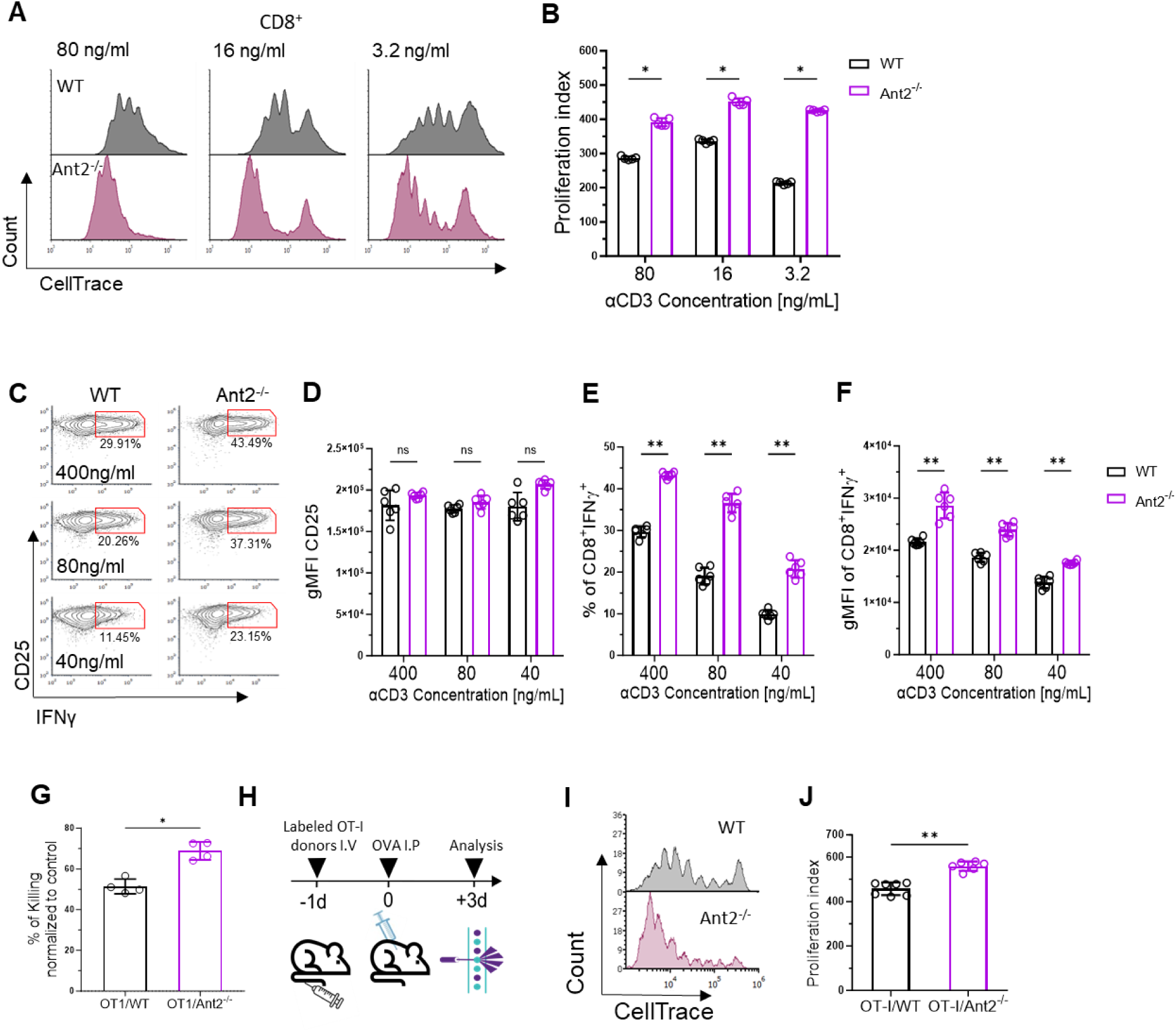
Increased activation potential and effector functions of Ant2^−/−^ CD8^+^ T cells. **(A)** Representative flow cytometry histograms of CellTrace intensity gated on CD8^+^ T cells derived from WT or Ant2^−/−^ mice, 72h post-activation with the indicated concentrations of αCD3 antibody and half the amount of αCD28 antibody. **(B)** Bar graph summarizing the results in A as a proliferation index. **(C)** Representative flow cytometry histograms showing intracellular IFNγ vs. CD25 gated on CD8^+^ T cells derived from WT or Ant2^−/−^ mice, 24h post-activation with the indicated concentrations of αCD3 antibody and half amount of αCD28. **(D-F)** Bar graphs summarizing the results shown in C, as geometric mean fluorescence intensity (gMFI) of CD25 **(D)**, the frequency of CD8^+^ IFNγ^+^ T cells **(E)**, and gMFI of IFNγ gated on CD8^+^IFNγ^+^CD25^+^ **(F)**. **(G)** WT or Ant2^−/−^ OT-I CD8^+^ T cells were activated with αCD3/αCD28 antibodies for 5 days. Then, cells were incubated with either B16 (control) or B16 melanoma-expressing ovalbumin target cells (B16-OVA) in a 1:1 ratio. Six hours later, the cells were washed three times with PBS. The remaining cells were then subjected to an MTT assay. **(H)** Schematic of the adoptive transfer experiment testing activation of naïve CD8^+^ T cells. CD45.1^+/+^ WT recipient mice were intravenously administered CellTrace-labeled CD45.2^+/+^ T cells derived from WT or Ant2^−/−^ OT-1 mice. After one day, recipient mice were intraperitoneally injected with 50µg of OVA. On day three following cell transfer splenocytes from the mice were examined by flow cytometry for CellTrace intensity. **(I)** Representative flow cytometry histograms of CellTrace intensity gated on donor (CD45.2^+/+^) CD8^+^ T cells. **(J)** Graph depicting the proliferation index of donor WT and Ant2^−/−^ OT-I T cells. Statistical method: Two-tailed Mann–Whitney test. Data are represented as mean ± S.D. (P value, *P≤0.05, **P≤0.01).

In an effort to investigate peptide-specific T-cell responses, we generated OT-I/Ant2^−/−^ mice and compared their responses to those of OT-I/WT littermates.^21^ Noteworthy, the OT-I model did not ameliorate the lymphopenia resulting from Ant2 deficiency (Figure S3A-B). To compare peptide-specific response while minimizing potential lymphopenia-related effects, we conducted a co-culture experiment. To distinguish between T cell populations, CFSE-labeled OT-I/WT and CellTrace-labeled OT-I/Ant2^−/−^ splenocytes were co-cultured in approximately a 1:1 ratio of CD8^+^ T cells (Figure S3C). These co-cultured cells were subsequently activated with OVA-peptide (SIINFEKL) for 24 hours. Our findings replicated our prior results using antibody activators and demonstrated that OT-I/Ant2^−/−^ CD8^+^ T cells exhibited significantly higher percentages of CD25^+^IFNγ^+^ cells and increased IFNγ intensity per cell (Figure S3D-E) compared to their WT counterparts. To further examine the effector function, we assessed the cytotoxic activity of OT-I/Ant2^−/−^ CD8^+^ effector T cells in an *in-vitro* killing assay.^22^ B16 melanoma cells expressing OVA (B16-OVA) and control cells (B16) were incubated with either OT-I/WT or OT-I/Ant2^−/−^ CD8^+^ effector T cells for 6 hours. The survival of B16-OVA cells incubated with Ant2^−/−^ derived CD8^+^ effector T cells was significantly lower compared to WT cells (Figure 3G). These results highlight the enhanced cytotoxic capacity of Ant2^−/−^ CD8^+^ T cells in an *in-vitro* setting, suggesting their enhanced ability to eliminate target cells.

Finally, we conducted an *in-vivo* experiment to evaluate the activation of OT-I/Ant2^−/−^ CD8^+^ T cells. CellTrace-labeled CD8^+^ T cells derived from CD45.2 OT-I/WT or OT-I/Ant2^−/−^ mice were intravenously administered to CD45.1 WT recipient mice. After one day, the recipient mice were intraperitoneally injected with 30µg of OVA. On day three following cell transfer, splenocytes from the recipient mice were collected and analyzed for CellTrace intensity. Consistent with our *in-vitro* findings, Ant2^−/−^ CD8^+^ T cells exhibited an increased proliferation *in-vivo* (Figure 3I-J).

In summary, our results demonstrate that naïve Ant2^−/−^ CD8^+^ T cells, in contrast to WT cells, exhibit decreased proliferation capacity during homeostatic expansion. Concurrently, they display an elevated sensitivity to activation signals, alongside improved proliferation capabilities and enhanced effector functions.

### Increased mitochondria biomass and spare respiratory capacity in Ant2^−/−^ T cells

The mitochondria play a central role in cellular metabolism and energy production, making them vital for T-cell activation and effector functions.^23,24^ Considering its pivotal role in the ADP/ATP exchange between the mitochondria and the cytosol, Ant2 deficiency leads to a decrease in mitochondria-originated ATP levels in the cytosol. Simultaneously, it causes a reduction in ADP levels in the mitochondrial matrix. Since ADP concentration drives ATP-synthase complex activity, its decrease reduces this activity, triggering proton accumulation in the intermembrane space. Consequently, this leads to increase ΔΨ, ultimately culminating in an inhibition of oxidative phosphorylation (OXPHOS).^25,26^ Indeed, naïve Ant2^−/−^ CD8^+^ and CD4^+^ T cells displayed substantially increased ΔΨ, as shown by Tetramethylrhodamine, methyl ester (TMRM) staining (Figure 4A-B and S4A-B).

**Figure 4:**
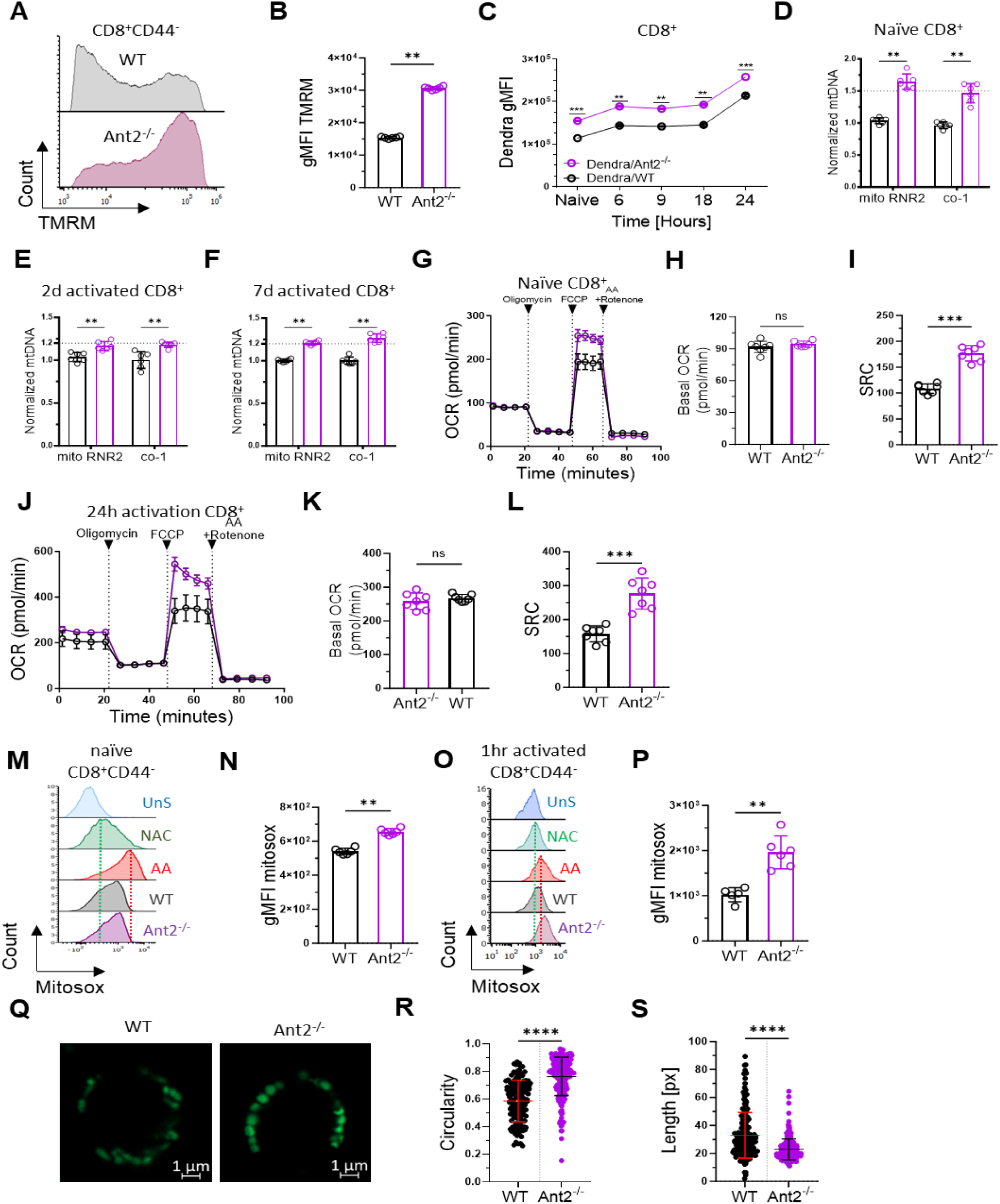
Increased biomass, and altered function, of mitochondria in Ant2^−/−^ T cells. **(A)** Representative flow cytometry histogram of Tetramethylrhodamine, Methyl Ester, Perchlorate (TMRM) staining gated on naïve (CD44^−^) CD8^+^ T cells. **(B)** Bar graph summarizing the results shown in A. **(C)** Dendra2 intensity of naïve and activated CD8^+^ T cells derived from WT and Ant2^−/−^ mito-Dendra2 littermate mice at the indicated time points. **(D-F)** Relative expression of Mitochondrial DNA (mtDNA) in naïve **(D)**, activated for 2d **(E)** or activated for 7d **(F)** WT or Ant2^−/−^ CD8^+^ T cells, measured by RT-PCR using 2 different set of primers targeted to amplify two mitochondrial genes, RNR2 or co-1. Results were normalized to the expression of the nuclear gene, β2M. **(G)** Continuous dot graphs of oxygen consumption rate (OCR) measured by Seahorse XF96, following consecutive injections of Oligomycin [1 µM], FCCP [2 µM], Rotenone + Antimycin-A [1 µM] (Rot+AA) of isolated naïve CD8^+^ T cell from WT (black) or Ant2^−/−^ (purple) mice at the indicated time points. **(H, I)** Bar graphs summarizing the results shown in G, as basal OCR **(H)** and spare respiratory capacity (SRC) calculated as the maximal OCR (after FCCP injection) minus Basal OCR **(I)**. **(J)** Same as in G using activated CD8^+^ T cells for 24h. **(K, L)** Bar graphs summarizing the results shown in J, as basal OCR **(K),** and SRC **(L)**. **(M-P)** Representative flow cytometry histograms of MitoSOX® staining gated on **(M)** unstimulated (naïve) and **(O)** one-hour activated CD8^+^ T cells. WT CD8^+^ T cells were used as controls, with the following conditions from top to bottom: unstained (UnS), pre- treated with 200µM N-Acetyl Cysteine (NAC), and treated with 1µM Antimycin A (AA). **(N, P)** Bar graphs summarizing the results of **(M)** unstimulated and **(O)** stimulated CD8^+^CD44^−^ T cells as gMFI of MitoSOX®. **(Q- S)** Representative live imaging of mitochondrial morphology in purified CD8^+^ T cells derived from WT and Ant2^−/−^ mito-Dendra2 littermate mice. High-resolution Images were captured using Zeiss laser scanning microscopy (LSM) 880 Airyscan, equipped with a 63X oil objective (3X zoom). The fluorescence of samples was excited at 425 nm. Scale bars are provided in the figures. **(R-S)** Quantitative analysis of the images was performed using NIS-Elements software (Nikon), allowing for the specific measurements of mitochondria length **(R)** and circularity **(S)**. Over n=300 mitochondria were analyzed for each genotype. Statistical method: Two-tailed Mann– Whitney test. Data are represented as mean ± S.D (B-F, H, I, K-S), or data are represented as mean ± SEM (G and J). (P value, **P≤0.01, ***P≤0.001, ****P<0.0001).

To assess the impact of Ant2 deficiency on mitochondrial biomass, we crossbred Ant2^−/−^ mice with those carrying the mitochondria-specific fluorescent protein Dendra2 (mtDendra2).^27^ mtDendra2 protein is expressed in various tissues, including T cells, allowing live-cell imaging, and enabling a direct assessment of mitochondrial biomass dynamics. By utilizing flow cytometry, we quantified the intensity of mtDendra2 in CD8^+^ T cells during various stages of *in-vitro* activation. Naïve and activated Ant2^−/−^ CD8^+^ T cells showed increased intensity per cell compared to WT cells, indicating higher mitochondrial content (Figure 4C).

In addition, to corroborate the association between the increase in mitochondrial biomass and Cre-mediated Ant2 deletion, we assessed mtDenra2 intensity in thymocytes across different developmental stages, both pre- and post-cre-mediated deletion, as well as in mature T cells within the spleen. Cre-mediated deletion by the distal Lck-promoter takes place as early as in the TCRβ selection stage (TCRβ^int^), reaching its peak at the TCRβ^hi^ stage.^28^ Intriguingly, when comparing the ratio of mtDenra2 intensity between Ant2^−/−^ and WT T cells at each developmental stage, a notable surge in mitochondrial biomass was observed precisely at the TCRβ^hi^ stage. This augmentation was further amplified in splenic CD8^+^ T cells within the Ant2^−/−^ mice context (Figure S4C-D). Collectively, these results support the notion that the loss of Ant2 is correlated with a significant increase in mitochondrial biomass over time. Furthermore, we evaluated mitochondrial DNA copy number using RT-PCR by comparing the ratio of two target mitochondrial genes (*MT-RNR2* and *MT-CO1*) to a reference nuclear gene (β2M). Naïve and activated Ant2^−/−^ CD8^+^ T cells exhibited higher mitochondrial DNA copy number compared to WT cells (Figure 4D-F). Together, these findings strongly suggest that Ant2 loss in naïve T-cell results in increased mitochondrial content.

We then examined the effect of Ant2 deficiency on the mitochondrial function of CD8^+^ T cells.

Using Seahorse analysis,^31^ we compared the oxygen consumption rate (OCR) of naïve and activated CD8^+^ T cells from both WT and Ant2^−/−^ mice. ^29^ The basal OCR of naïve and activated Ant2^−/−^ CD8^+^ T cells was similar to that of their WT counterparts (Figure 4G-H, J-K). Considering the increased mitochondrial content in Ant2^−/−^ T cells, our data imply that their basal OCR per mitochondrion is diminished compared to WT controls. Interestingly, when treated with the uncoupler FCCP, which dissipates membrane hyperpolarization, naïve and activated Ant2^−/−^ CD8^+^ T cells exhibited a significant increase in maximal OCR compared to WT cells, resulting in an increased spare respiratory capacity (SRC) (Figure 4I, L).^30^ Thus, we propose that the increase in mitochondrial biomass is a result of Ant2 deficiency, serving as a compensatory response to OXPHOS inhibition arising from decreased mitochondrial ADP levels and persistent membrane hyperpolarization.

The cellular antioxidant defense mechanisms tightly regulate mitochondrial ROS generation.^6^ However, OXPHOS inhibition imposed by the loss of Ant2 may disturb this equilibrium. Indeed, MitoSOX® staining revealed that during the naïve state, Ant2^−/−^ CD8^+^ T cells display a modest, yet significant increase in ROS production compared to their WT counterparts. Upon one hour of stimulation, this imbalance becomes even more pronounced, as both Ant2^−/−^ CD4^+^ and CD8^+^ T cells exhibit a further amplification of ROS production (Figure 4M-P, S4E-F). Consistent with these findings, our high-resolution live-cell microscopy revealed that the mitochondria of Ant2^−/−^ CD8^+^ T cells exhibited significantly higher circularity scores and lower length scores compared to WT cells (Figure 4Q-S), indicative of a blob-shaped mitochondrial morphology associated with the accumulation of ROS within the mitochondria.^31,32^ Altogether, our results support the concept that Ant2 deficiency leads to a reduction in ADP concentration within the mitochondrial matrix, ultimately resulting in insufficient cellular respiration, decreased OXPHOS, an inherent increase in ROS generation, and alterations in mitochondrial morphology. In response to this respiratory bottleneck, Ant2^−/−^ T cells increase their mitochondrial content and rewire mitochondrial function.

### Primed-like proteome and distinct mitochondria in naïve Ant2^−/−^ CD8^+^ T cells

To gain insights into the metabolic changes associated with Ant2 deficiency in naïve CD8^+^ T cells, we employed protein mass spectrometry (MS) to examine alterations in the cellular proteome. Comparative analysis revealed that the protein expression profiles of naïve Ant2^−/−^ CD8^+^ T cells were largely similar to WT cells (Figure S5A), although we observed 412 upregulated and 610 downregulated proteins in Ant2^−/−^ compared to WT cells (Figure S5B). Focusing on the 535 mitochondrial-associated proteins detected in our experiment,^33^ we identified 56 enriched proteins in Ant2^−/−^ cells vs. 15 in WT (Figure 5A, proteins marked in red). Of note, consistent with the findings of unaltered Ant1 mRNA expression levels (Figure 1C), there was no significant change observed in Ant1 protein expression levels (Figure 5A).

**Figure 5:**
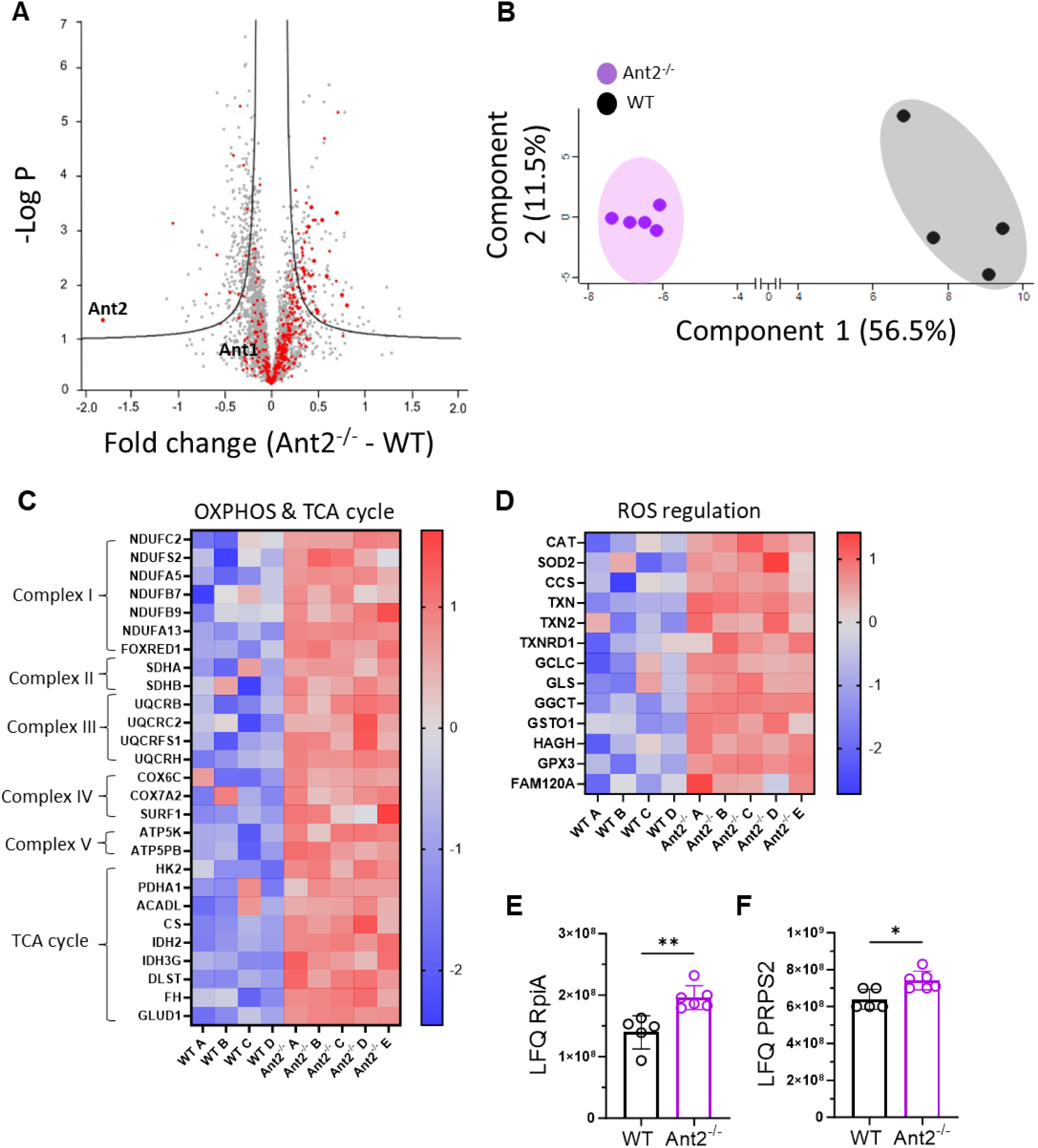
Altered proteome and distinct mitochondria in naïve Ant2^−/−^ CD8^+^ T cells. **(A-F)** Mass-spectrometry analysis was performed on protein extracts from naïve (CD44^−^) CD8^+^ T cells derived from WT or Ant2^−/−^ OT-I mice (n=4 or 5 in each group). **(A)** Volcano plot displaying differentially expressed proteins (relative peptide abundances). The negative logarithm of statistical significance (*p*-*value*) is plotted against the fold change magnitude. Prior to imputation from a normal distribution, Label-Free Quantitation (LFQ) levels were log_2_- transformed. Red dots represent mitochondrial-associated proteins sourced from MitoCarta2.0. **(B)** Principal component analysis comparing the log_2_-transformed data of all mitochondrial-associated proteins. The first two principal components, which capture the highest variance in the data, are depicted. **(C-D)** Heatmaps illustrating differentially expressed proteins associated with oxidative phosphorylation & tricarboxylic acid cycle **(C)**, as well as reactive oxygen species regulation & glutathione synthesis **(D)**. The heatmaps were generated based on the log2 fold change values of individual genes between the two compared samples. The color gradient ranges from red, indicating a significant fold change, to blue, representing a minor fold change. **(E-F)** Bar graphs displaying the LFQ levels of RpiA **(E)** and Prp2 **(F)**. Statistical method: Two-tailed multiple t-tests, FDR <0.05, S0<0.1(A), or by Benjamini-Hochberg method (FDR <0.05) (B), or by student’s t-test analysis conducted on log2-transformed data after the Z-score normalization step (C and D), or by two-tailed Mann–Whitney test (E and F). Data are represented as mean ± S.D. (P value, *P≤0.05, **P≤0.01).

Further, in line with the observed increase in mitochondria biomass, our proteomic analysis revealed an elevated expression of the voltage-dependent anion-selective channel protein 1 (VDAC1) (Figure S5C). Moreover, we observed the upregulation of key mitochondrial proteins associated with mitochondrial biogenesis in naïve Ant2^−/−^ CD8^+^ T cells. These proteins include transcription factor A (TFAM) and deoxyguanosine kinase (DGUOK), both play a critical role in mitochondrial replication.^34,35^ In addition, proteome-derived principal component analysis (PCA) comparing all detected mitochondria-associated proteins in both samples, revealed that in addition to increased mitochondrial content, Ant2^−/−^ T cells hold distinct mitochondria in comparison to WT cells (Figure 5B).

An in-depth analysis of the proteomic data unveiled several upregulated metabolic pathways in Ant2^−/−^ CD8^+^ T cells, both in mitochondrial and cytosolic compartments. The most significant upregulated pathways were OXPHOS-related proteins (such as NDUFB9, SDH, UQCRFS1, COX6C, and ATP5K, which constitute representatively proteins of complex (C)I, CII, CIII, CIV, and CV, respectively) and TCA cycle-related proteins (including CS, IDH2, and FH) (Figure 5C). These pathways play critical roles not only in adapting to insufficient OXPHOS but also in supporting T-cell activation and effector functions.^24,36^ In addition, our MS analysis revealed the upregulation of crucial detoxification mechanisms. The observed adaptation resembles that of activated T cells and involves the upregulation of key antioxidant enzymes such as SOD2, catalase, and thioredoxin, along with the enhancement of the glutathione system (Figure 5D).^37,38^ These results are in line with our increased MitoSOX® staining in Ant2^−/−^ T cells and suggest an attempt to maintain redox homeostasis. The activity of this anti-ROS machinery and the maintenance of reduced glutathione is dependent on the availability of NADPH, suggesting the involvement of pathways for NADPH generation. Indeed, our proteomic analysis further revealed the enrichment of enzymes associated with the PPP (Figure 5E-F), supporting the hypothesis that constant NADPH consumption stimulates G6PD activity.^39^ Additionally, the proteomic analysis uncovered alterations in proteins related to folate-dependent one-carbon metabolism (Figure S5D),^9,40,41^ iron and heme metabolism (Figure S5E),^42–45^ and other key enzymes that are typically upregulated upon activation. These enzymes are discussed in more detail in the next paragraph. Pathway enrichment analysis conducted on all upregulated proteins in Ant2^−/−^ T cells using Metascape,^46^ verified a significant enrichment of pathways primarily associated with mitochondrial functions (Figure S5F). These findings support the notion that the loss of Ant2 leads to both increased mitogenesis and mitochondria rewiring in naïve CD8^+^ T cells.

Overall, our proteomic analysis provides valuable insights into the metabolic adaptations occurring in Ant2^−/−^ naïve T cells. These results further corroborate our notion that naïve Ant2-deficient CD8^+^ T cells acquire distinct mitochondria and a primed-like proteome, which contributes to the increased propensity toward activation.

### Specialized mitochondria and activated-like metabolic signature in naïve Ant2^−/−^ CD8^+^ T cells

Next, we postulated that the observed alterations in the proteome and mitochondrial reprogramming in naïve T cells lacking Ant2 would result in a rewired glucose metabolism, ultimately contributing to the enhanced phenotype upon activation. Therefore, we cultured OT-I/WT and OT-I/Ant2^−/−^ naïve CD8^+^ T cells with ^13^C_6-_glucose for four hours and analyzed both the cellular and media fractions using liquid chromatography-mass spectrometry (LC-MS). This approach allowed us to assess the incorporation of ^13^C into central metabolites and provide a comprehensive understanding of the alterations in glucose metabolism resulting from Ant2 deficiency.

Naïve Ant2^−/−^ CD8^+^ T cells exhibited a marked decrease in unlabeled phosphocreatine (PCr) to creatine levels (Figures 6A, S6A-B). In line with this result, our proteomic data indicate elevated Creatine Kinase B (CKB) levels in naïve Ant2^−/−^ CD8^+^ T cells (Figure 6B). CKB, the primary isoform of creatine kinase in CD8^+^ T cells,^47^ catalyzes the phosphoryl transfer from ATP to creatine and vice versa. Previous studies have demonstrated that CKB is crucial for ATP buffering in activated T cells and other cells that rapidly consume ATP.^47,48^ The creatine-PCr shuttle was also proposed to serve as an energy transport between subcellular sites of ATP production.^49^ Consequently, we propose that the high ATP level within the mitochondrial matrix of Ant2^−/−^ T cells stimulate the conversion of creatine to Pcr. Pcr is then transported to the cytosol and used by CKB to phosphorylate ADP to ATP, forming a circuit that enables Ant2^−/−^ T cells to compensate for the lack of direct mtATP transfer.

**Figure 6:**
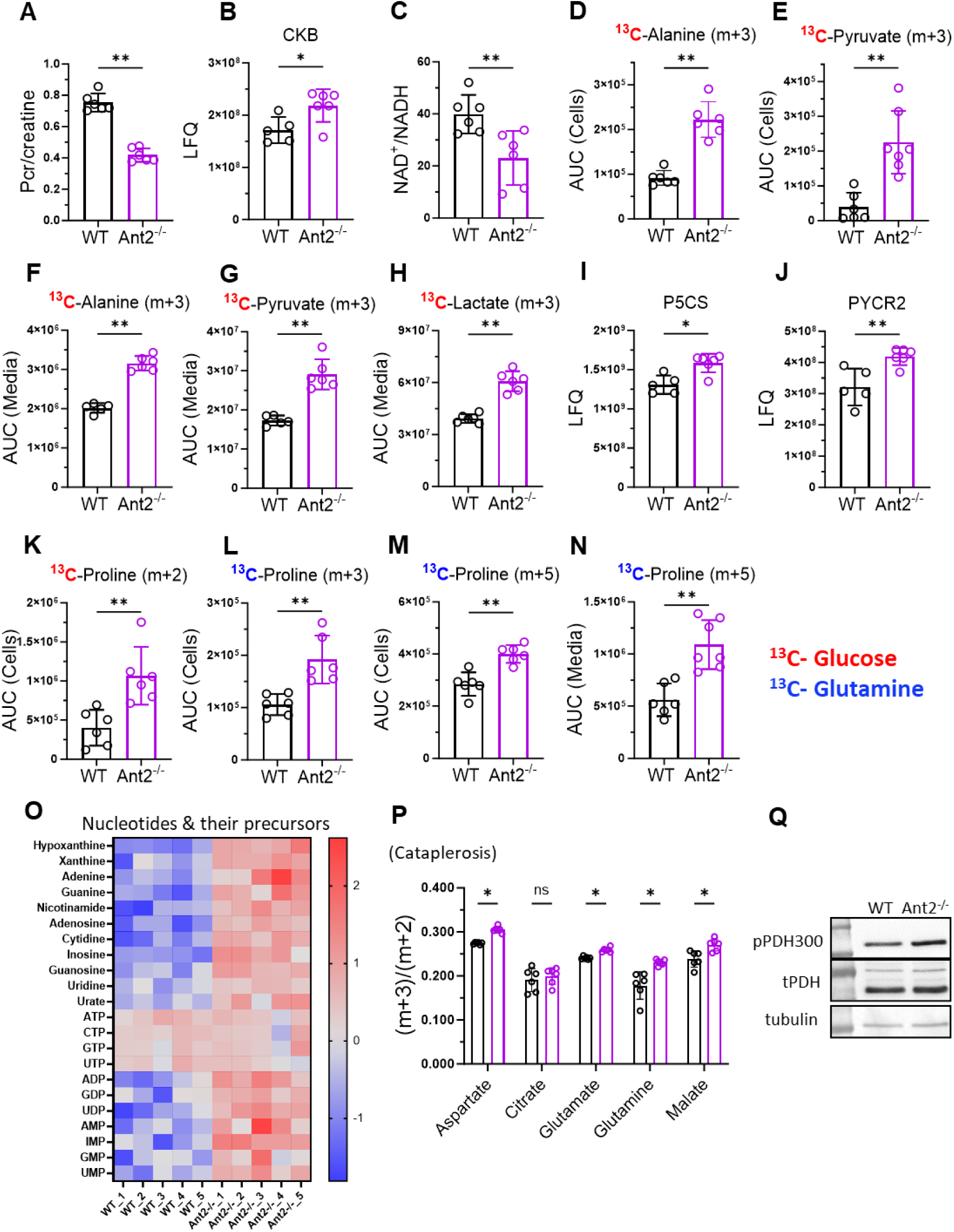
Metabolic adaptations of naïve Ant2^−/−^ CD8^+^ T cells. **(A-O)** Comparison of the area under the curve (AUC) values obtained from metabolomics analysis combined with ^13^C_6_-glucose or ^13^C_5_-glutamine tracing, and the Label-Free Quantitation (LFQ) values attained from proteomics analysis, between naïve (CD44^−^) WT and Ant2^−/−^ CD8^+^ T cells. **(A)** Bar graph showing the ratio between the AUC values of unlabelled Phosphocreatine (Pcr) and Creatinine. **(B)** Bar graph displaying the LFQ values of Creatine Kinase. **(C)** Bar graphs demonstrating the NAD^+^/NADH ratio. **(D-H)** LC-MS measurements of labeling patterns for glycolytic derivatives following [^13^C6]-glucose tracing. Bar graphs showing the AUC values of alanine m+3 **(D, F)**, and pyruvate m+3 **(E, G)** in the cell and media fractions, as well as lactate m+3 in the media fraction **(H)**. **(I-J)** Bar graphs illustrating the LFQ values of the two crucial mitochondrial enzymes involved in the proline biosynthesis pathway, P5CS **(I)** and PYCR2 **(J)**. **(K-N)** Bar graphs displaying AUC from LC-MS measurements of proline labeling patterns via ^13^C6- glucose tracing (m+2) in the cell fraction **(K)**, and ^13^C5-glutamine tracing (m+3 and m+5) in the cell fraction **(L, M)**, as well as m+5 in the media fraction **(N)**. **(O)** The Heatmap illustrating differentially expressed unlabeled nucleotides and their precursors, detected through our carbon tracing experiments. The heatmap was generated based on the log2 fold change of individual metabolites between the two samples. The color gradient spans from red, indicating a substantial fold change, to blue, representing a minimal fold change. **(P)** Bar graphs illustrating the (m+3) to (m+2) ratio of specific TCA cycle intermediates and TCA cycle-derived metabolites in naïve (CD44^−^) WT and Ant2^−/−^ CD8^+^ T cells. The metabolites analyzed include (left to right) aspartate, citrate, glutamate, glutamine, and malate. **(Q)** Representative immunoblot analysis of whole-protein extracts from purified WT and Ant2^−/−^ CD8^+^ cells using anti-total PDH (tPDH) and pPDH^ser300^ antibodies. An anti-tubulin antibody was used as a loading control. Statistical method: two-tailed Mann–Whitney test (A-N, P), or by student’s t-test analysis conducted on log2-transformed data after the Z-score normalization step (O). Data represented as the mean ± S.D. (P value, *P≤0.05, **P≤0.01).

Naïve Ant2^−/−^ CD8^+^ T cells also displayed a decreased NAD^+^/NADH ratio (Figure 6C), indicating altered redox balance. In addition, we observed a significant increase in both cellular and media levels of labeled pyruvate m+3 and alanine m+3, as well as lactate m+3 secretion (Figure 6D-H) compared to the control group. These findings suggest a rewiring of pyruvate metabolism in Ant2^−/−^ T cells due to OXPHOS inhibition.^50^ Notably, this metabolic signature shares similarities with activated T cells whereas the demand for NAD^+^ exceeds OXPHOS capacities.^10^

The impermeability of the inner mitochondrial membrane to NAD(P)H presents a challenge in transferring reducing equivalents to the mitochondria. Consequently, mitochondria have evolved various shuttles and para-metabolic pathways to regenerate NAD^+^ pools. Proline is a non-essential amino acid that is derived from glutamate.^51^ Other than being required for efficient protein biosynthesis post-activation, it did not receive high attention in the context of T cells. Recently, evidence resurfaced that proline biogenesis may serve as a para-metabolic pathway for regenerating mitochondrial NAD^+^ in highly proliferative cancerous cells, where OXPHOS is insufficient to regenerate adequate NAD^+^ levels.^51–55^

In the context of naïve Ant2^−/−^ CD8^+^ T cells, our MS-proteomics analysis revealed the upregulation of the enzymes involved in proline biosynthesis, pyrroline-5-carboxylate synthase (P5CS) and mitochondrial pyrroline-5-carboxylate reductase (PYCR2) (Figure 6I-J). These enzymes catalyze the reduction of glutamate to pyrroline-5-carboxylate (P5C), which is further reduced to proline. Interestingly, our ^13^C_6_ glucose-tracing experiments showed an increase in labeled proline m+2 in naïve Ant2^−/−^ CD8^+^ T cells (Figure 6K), indicating elevated proline biosynthesis even in the naïve state.

We further investigated the source of glutamate for proline biosynthesis using ^13^C_5_-glutamine tracer analysis. As expected, proline m+3 and proline m+5 were elevated in Ant2^−/−^ CD8^+^ T cells, suggesting enhanced proline biosynthesis from glutamine as a substrate (Figure 6L-M). Furthermore, a significant portion of the newly synthesized proline m+5 was secreted from the cells (Figure 6N). Considering its secretion, and the fact that naïve CD8^+^ T cells are catabolic in nature, it is unlikely that the synthesized proline is solely for protein synthesis purposes. Rather, we propose that Ant2^−/−^ naïve CD8^+^ T cells engage in proline biosynthesis as a para-metabolic pathway to regenerate mitochondrial NAD(P)^+^ in an attempt to compensate for the reduced OXPHOS. This metabolic adaptation likely contributes to maintaining redox balance and increased aerobic glycolysis and the PPP in Ant2^−/−^ T cells.^56^ Notably, our glucose-tracing analysis provides additional support for the increased PPP activity in Ant2^−/−^ CD8^+^ T cells, as indicated by elevated levels of unlabeled nucleotides and their precursors (Figure 6O). Specific labeled metabolites, such as inosine m+5 and ATP m+5 provide additional indications of increased nucleotide metabolism in Ant2^−/−^ CD8^+^ T cells (Figure S6 C-D).

Our findings thus far suggest that naïve Ant2^−/−^ CD8^+^ T cells have successfully rewired their mitochondrial metabolism to overcome the constrained OXPHOS. Indeed, we observed no signs of TCA cycle congestion in these cells, as the labeling patterns of TCA cycle intermediate metabolites remained comparable to WT cells (Figure S6E-H). In fact, we observed upregulation of labeled TCA cycle-derived amino acids (Figure 6K-L, Figure S6I-K), indicating an enhanced cataplerosis.^36,57^

Cataplerosis, the process of carbon exiting the TCA cycle for precursor generation, is known to occur during T-cell stimulation, enabling the cells to generate macromolecules for proliferation.^58^ Given the observed increase in carbon efflux in naïve Ant2^−/−^ CD8^+^ T cells, there must be a corresponding increase in carbon influx, termed anaplerosis, to compensate for the lost carbon.^36^ Our glutamine tracing experiments exclude glutamine as an anaplerotic source (data not shown), which is expected as glutamine addiction occurs upon activation. Thus, we explored an alternative anaplerosis source.

Pyruvate carboxylase (PC) converts pyruvate to oxaloacetate, replenishing the TCA cycle with a 4-carbon input instead of the typical 2-carbon input (as acetyl-CoA), mediated by pyruvate dehydrogenase (PDH). PC activity is critical for activated T-cell proliferation and eliciting anti-tumor responses *in-vivo*.^59,60^ While direct measurement of PC activity is challenging, indirect activity can be inferred from certain conditions and measurements. First, we anticipated increased PC activity due to inhibited OXPHOS and increased ROS production, as previous studies have demonstrated their correlation.^61^ Secondly, leveraging our ^13^C_6_-glucose tracing data, we compared the m+3 to m+2 ratios of TCA cycle derivatives as a proxy for PC vs. PDH activities. M+3 indicates direct pyruvate-derived oxaloacetate contributed by PC, while m+2 indicates pyruvate-derived acetyl-CoA by PDH. Our analysis indeed revealed elevated m+3/m+2 isotopologue ratios of aspartate, glutamate, and glutamine in Ant2^−/−^ compared to WT, suggesting a relative increase in PC contribution (Figure 6P). Third, we examined the inhibitory phosphorylation status of PDH as an indication for PC activity.^60,62^ Immunoblot analysis revealed an increase in phosphorylated PDH on serine 300 in naïve Ant2^−/−^ CD8^+^ T cells compared to WT cells (Figure 6Q), suggesting increased pyruvate flow towards other available pathways, such as PC and lactate dehydrogenase (LDH).

Lastly, we observed a prominent glucose-derived aspartate (m+2 and m+4) secretion in naïve Ant2^−/−^ CD8^+^ T cells (Figure S6L). While aspartate is crucial for nucleotide biosynthesis in activated T cells, the same demand is not typical of naïve T cells. The marked synthesis of aspartate in Ant2^−/−^ naïve cells may explain the increased levels of nucleotides and their precursors in these cells. Additionally, it hints at the potential alteration in carbon efflux dynamics, with these cells diverting excess carbon away from the TCA cycle due to increased PC activity in naïve state. Notably, in effector cells, increased PC activity leads to succinate secretion as a 4-carbon output.^60^ Nevertheless, it appears that increased PC activity serves as a metabolic adaptation to redirect excess carbons toward anabolic pathways and effector functions.

Importantly, we conducted a series of glucose concentration-dependent experiments to exclude the possibility of increased glucose dependency or uptake. *In-vitro* assays for IFNγ production (Figure S7A-D) and proliferation (Figure S7E-G) were performed, along with glucose uptake studies using 2-NBDG Glucose (Figure S7H-I). We found no evidence of increased glucose dependency or uptake. Instead, these results strongly suggest that the observed metabolic alterations in Ant2-deficient T cells are primarily attributed to altered glucose utilization.

Overall, our findings highlight the metabolic adaptations enforced by Ant2 deficiency in T cells, which include aerobic glycolysis, TCA cycle activity, nucleotide metabolism, increased ROS production, and proline biosynthesis. These metabolic alterations collectively contribute to the anabolic changes observed in Ant2-deficient T cells, resulting in improved effector function, increased proliferation, and the ability to cope with the expected energetic challenges.

### Pharmacological inhibition of ANT augments naïve T cell activation and anti-tumor activity

The results of our study demonstrate that disrupting mtATP transfer to the cytosol in naive T cells, achieved through Ant2 knockout, initiates metabolic rewiring that increases their propensity for activation. Therefore, we aimed to explore the therapeutic potential of these findings by using pharmacological pan-ANT inhibitors.^63^

Previous studies have shown that prolonged ANT inhibition by atractyloside (ATR) and carboxyatractyloside (CATR) *in-vivo* leads to metabolic rewiring, resulting in improved outcomes for the organism.^64–66^ We investigated the effects of prolonged ANT inhibition using both inhibitors on T-cell function *in-vivo*. To test T-cell activation, donor OT-I mice received daily injections of CATR (2.5 mg/kg) or PBS (control) for two weeks. The OT-I donor cells were labeled with CellTrace and then adoptively transferred intravenously into WT recipient mice. After 24 hours, the recipient mice were injected intraperitoneally with OVA. Seventy-two hours after the OVA injection, CD8^+^ T cells from the spleen of recipient mice were analyzed for CellTrace intensity and intracellular IFNγ staining after a brief *ex-vivo* restimulation (Figure 7A). As anticipated, ANT inhibition by CATR resulted in a significant increase in T cell proliferation (Figure 7B-C) and IFNγ production (Figure 7D-E).

**Figure 7:**
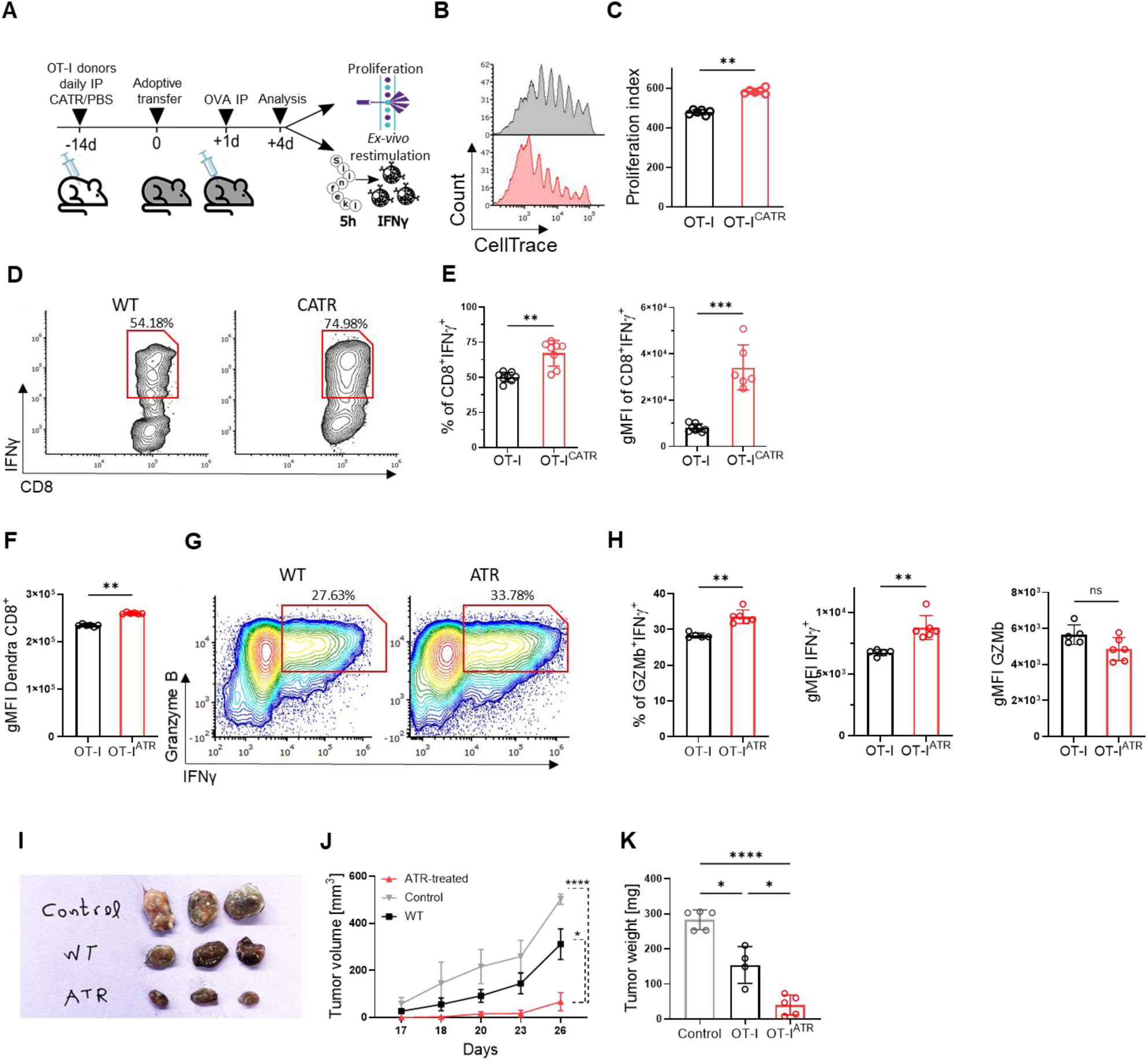
Pharmacologically Ant inhibition by atractyloside (ATR) and carboxyatractyloside (CATR) **Pharmacological inhibition of ANT heightens naïve T cell responsiveness to activation stimuli. (A)** Schematic of adoptive transfer experiment. OT-I/WT mice were intraperitoneally (i.p) administered daily with either 2.5mg/ml CATR or PBS for 14 days. After sacrificed, splenocytes were stained with CellTrace and adoptively transferred to WT recipient mice. One day after cell transfer, recipient mice received 100µg of OVA i.p. On the third day following cell transfer, splenocytes were analyzed by flow cytometry for CellTrace intensity and *ex-vivo* restimulation using SIINFEKEL peptide (1µg/ml) for 5 hours. **(B)** Representative flow cytometry histogram showing CellTrace intensity gated on donor PBS-treated (grey) or CATR-treated (red) WT OT-I T cells. **(C)** Graph depicting proliferation index of donor CATR-treated or PBS-treated WT OT-I T cells. **(D)** Representative flow cytometry plots showing CD8 vs. IFNγ staining gated on donor PBS-treated (left) or CATR-treated (right) WT OT-I T cells. **(E)** Bar graphs summarizing the results shown in D as the frequency of CD8^+^IFNγ^+^ T cells (left), and gMFI of IFNγ gated on CD8^+^IFNγ^+^ T cells (right). **(F-K)** Female mice were injected subcutaneously with 1 x 106 B16-OVA tumor cells. Mice were then divided into three groups (n=4 per group): control (no adoptive transfer), WT, and ATR-treated OT-I CD8+ T cell-injected. **(F)** Intensity of mtDendra2 in donor OT-I CD8+ T cells (both WT and ATR-treated), gated on naïve CD8+ T cells, immediately after spleen dissection. **(G)** Representative flow cytometry plots displaying IFNγ versus granzyme B (GzmB) staining in donor WT (left) and ATR-treated (right) CD8+ OT-I T cells. **(H)** Bar graphs summarizing the frequency of IFNγ+GzmB+ (left), gMFI of IFNγ+ cells (middle), and gMFI of GzmB+ cells (right), all gated on donor CD8+ OT-I T cells. **(I)** Images of tumors from control, WT, and ATR-treated OT-I CD8+ T cell-injected mice at the end of the experiment. **(J)** Tumor volume (mm³) was measured daily using calipers starting from day 17 post-tumor implantation. **(K)** Tumors were excised, weighed, and presented as tumor weight (mg). Statistical method: two-tailed Mann–Whitney test (C, E, F, H), or by two-way ANOVA test (J), or by one-way ANOVA test (K). Data are represented as mean ± S.D for all expect ±SEM (J). (P value, *P≤0.05, **P≤0.01, ***P≤0.001, ****P≤0.0001)

Next, to assess the effect of ANT inhibition on T cell anti-tumor activity and mitobiogenesis, we used the well-established B16-OVA tumor-bearing mouse model. Donor OT-I/mtDendra2 mice were pre-treated with either ATR (1mg/kg) or PBS (control) for 10 days. Subsequently, T cells from the spleen of the donor mice were immediately analyzed for mtDendra2 intensity before being activated *ex-vivo* with the SIINFEKEL peptide for 2 days. The activated T cells were then adoptively transferred into sub-lethally irradiated mice bearing B16-OVA tumors. Tumor growth was monitored daily. Our findings demonstrate that ATR-treated naïve T cells significantly increased mitochondrial biomass (Figure 7F), as indicated by mtDendra2 intensity. In addition, they also showed increased IFNγ production, with comparable granzyme b expression (Figure 7G-H). Furthermore, both OT-I and ATR-treated OT-I T cells inhibited tumor growth compared to control mice that did not receive adoptive cell therapy (ACT), with ATR-treated OT-I T cells showing significantly greater tumor growth inhibition compared to OT-I T cells alone (Figure 7I-K). Overall, these findings strongly support our overarching concept: restricting mtATP transfer to the cytosol leads to metabolic adaptations that enhance responsiveness to activation stimuli, which may be used for therapeutic purposes.

## Discussion

The immune system’s ability to respond effectively to threats requires a delicate balance between T-cell activation and regulation. Recent studies have unveiled the compelling role of metabolism in molding T-cell behavior, with metabolic reprogramming and checkpoints emerging as crucial determinants of T-cell fate and function.^5,24,67^ Indeed, these cells exhibit remarkable adaptability even in challenging circumstances and hostile surroundings. While other cell types might falter, T cells demonstrate an exceptional ability to adjust through diverse metabolic pathways, shuttles, and ‘venting’ strategies.^10,68^ Here, we explore the intricate interplay between mitochondrial dynamics, bioenergetics, and immune function by investigating the effects of indirect OXPHOS restriction induced by Ant2 knockout.

Ant2 deletion has varied tissue-specific effects. It protects against non-alcoholic fatty liver disease (NAFLD) in the liver, ^65^ improves glucose tolerance and insulin resistance in adipose tissue,^69^ and induces a proinflammatory phenotype in macrophages.^70^ Most recently, it protects against obesity-induced renal dysfunction.^66^ However, whole-body or cardiomyocyte-specific Ant2 knockout is lethal, emphasizing its critical role in these organs.^14,71^ Here, we explore the intricate interplay between mitochondrial dynamics, bioenergetics, and immune function by investigating the effects of indirect OXPHOS restriction induced by Ant2 knockout. To surmount this metabolic bottleneck, both the cytosol and mitochondria of Ant2^−/−^ T cells undergo significant metabolic adaptations. In an endeavor to restore the balance between NAD^+^ and NADH, naïve Ant2^−/−^ T cells increase mitochondrial biogenesis while concurrently engaging in aerobic glycolysis and proline biosynthesis. Additionally, the antioxidant defense mechanisms are upregulated to counter oxidative stress. Notably, ROS levels remain slightly elevated, which may explain the heightened responsiveness to activation stimuli.^72,73^ Further, the elevated NADP^+^/NADPH ratio originating from antioxidant activities together with the increased proline biosynthesis might prematurely induce glucose flux towards the PPP and subsequent nucleotide accumulation (Figure 8).^39^

**Figure 8:**
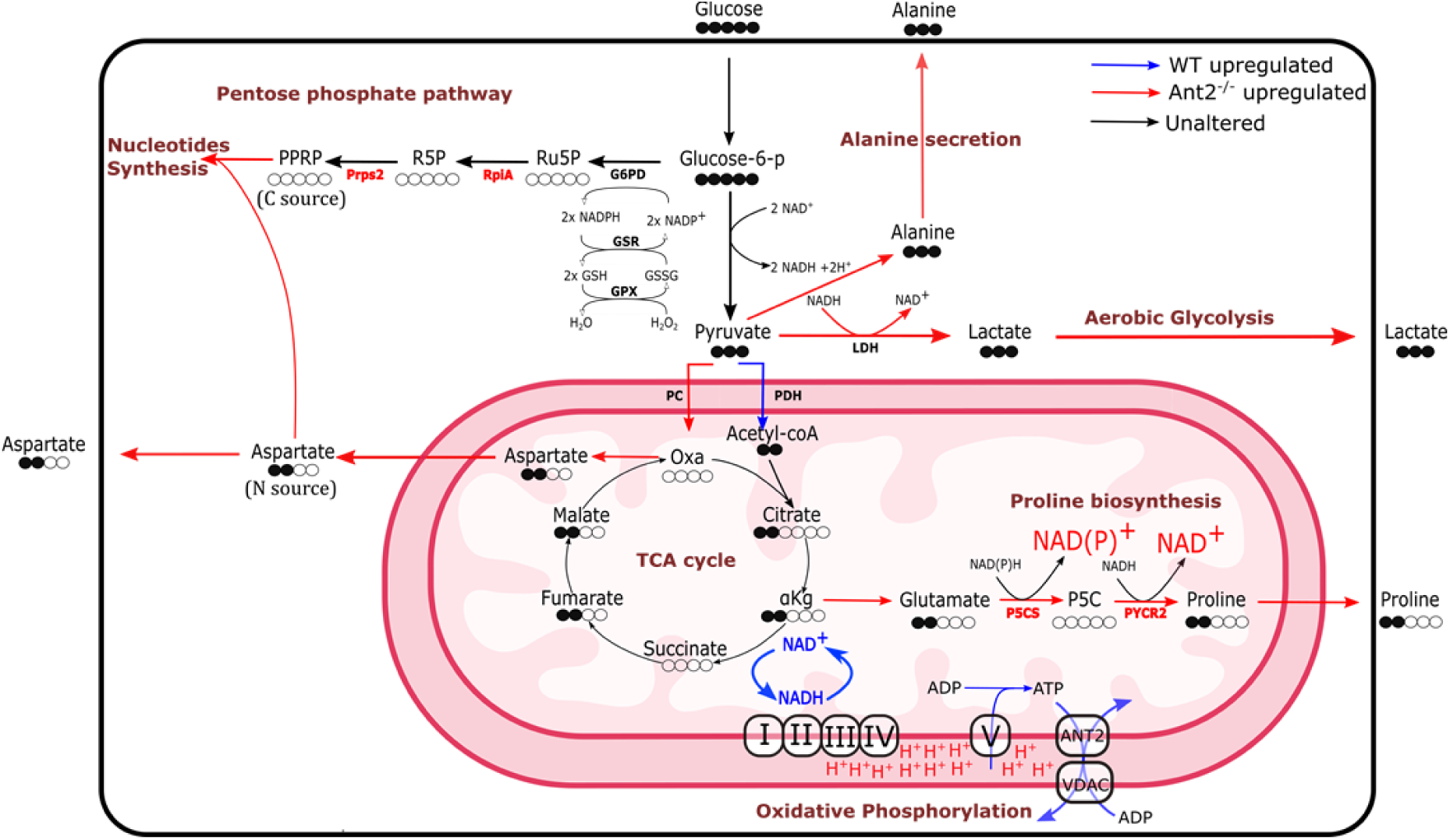
Summary of metabolic alterations imposed by Ant2-deficiency in naïve T cells. (1) Ant2 deficiency leads to a reduction in mtATP transfer to the cytosol, accompanied by a concomitant decrease in ADP concentrations within the mitochondrial matrix. (2) ATP-synthase complex functionality is hindered, resulting in mitochondrial membrane hyperpolarization and OXPHOS inhibition. (3) Increased aerobic glycolysis & excess carbon secretion as alanine and lactate. (4) Increased PC activity due to PDH inhibition. (5) Increased cataplerosis as TCA cycle derived amino-acids. (6) Increased anti-oxidative stress machinery. (7) Increased Phosphate pentose pathway.

Remarkably, many of these metabolic alterations observed in naive Ant2-deficient T cells echo the metabolic rewiring characteristic of recently activated T cells, albeit driven by a distinct rationale.^62,74^ In the context of activated T cells, the pursuit of biosynthetic precursors becomes paramount for enabling rapid proliferation.^75^ This heightened biosynthetic demand prompts increased OXPHOS activity, aimed at sustaining a favorable NAD^+^ to NADH ratio to support the oxidation reactions of the TCA cycle. OXPHOS upregulation gives rise to increased production of ROS, which further supports the activation signal.^72^ This orchestrated response also triggers the PPP, facilitating the synthesis of nucleotides crucial for rapid cell proliferation. Nonetheless, it is well-established that the demand for NAD^+^ outpaces that of ATP in rapidly proliferating cells,^10^ driving metabolic rewiring towards aerobic glycolysis and other alternative pathways for NAD^+^ regeneration, such as proline biosynthesis. Hence, we propose that OXPHOS inhibition and its ensuing cascade of events in Ant2-deficient naïve T cells prompt an adaptive response akin to the metabolic programming witnessed in primed cells. As a substantial portion of their metabolic rewiring has already occurred, these naïve T cells appear to bypass the typical metabolic reprogramming associated with priming. Consequently, these naïve T cells manifest activated-like metabolic characteristics and behaviors.

Building on the metabolic adaptations observed in Ant2-deficient naïve T cells, our results demonstrate that prolonged pharmacological ANT inhibition, using ATR and CATR, recapitulates our Ant2^−/−^ model in WT cells. Specifically, recipient mice receiving CATR-treated OT-I cells exhibited increased proliferation and IFNγ production upon restimulation, indicative of augmented activation. Similarly, in the B16-OVA tumor model, ATR-treated OT-I T cells demonstrated enhanced mitochondrial biomass, IFNγ production, and tumor growth inhibition compared to control T cells. These findings collectively suggest that restricting mtATP transfer to the cytosol can induce metabolic changes that potentiate T-cell activation. This approach holds promise for developing novel immunotherapies by leveraging ANT inhibitors to optimize T-cell function and efficacy against tumors, or perhaps even infectious diseases through targeted metabolic modulation.

Finally, our findings raise an intriguing question regarding the selective pressure for Ant2 function in T cells. Despite its plausible inhibitory effect on T cell activation, Ant2 is still selected to be expressed in T cells. Our results indicate that Ant2 deficiency prioritizes enhanced activation responses at the expense of homeostatic proliferation. Given that the balance between activation and homeostasis is crucial for immune system function, one can appreciate the potential role of Ant2 in upholding proper immunoregulation and averting overactivation and autoimmunity. Nonetheless, targeting Ant2 could hold beneficial effects in augmenting the effectiveness of therapeutic methods, like adoptive T cell transfer.

## Supplementary figures

**Supplementary Figure 1:**
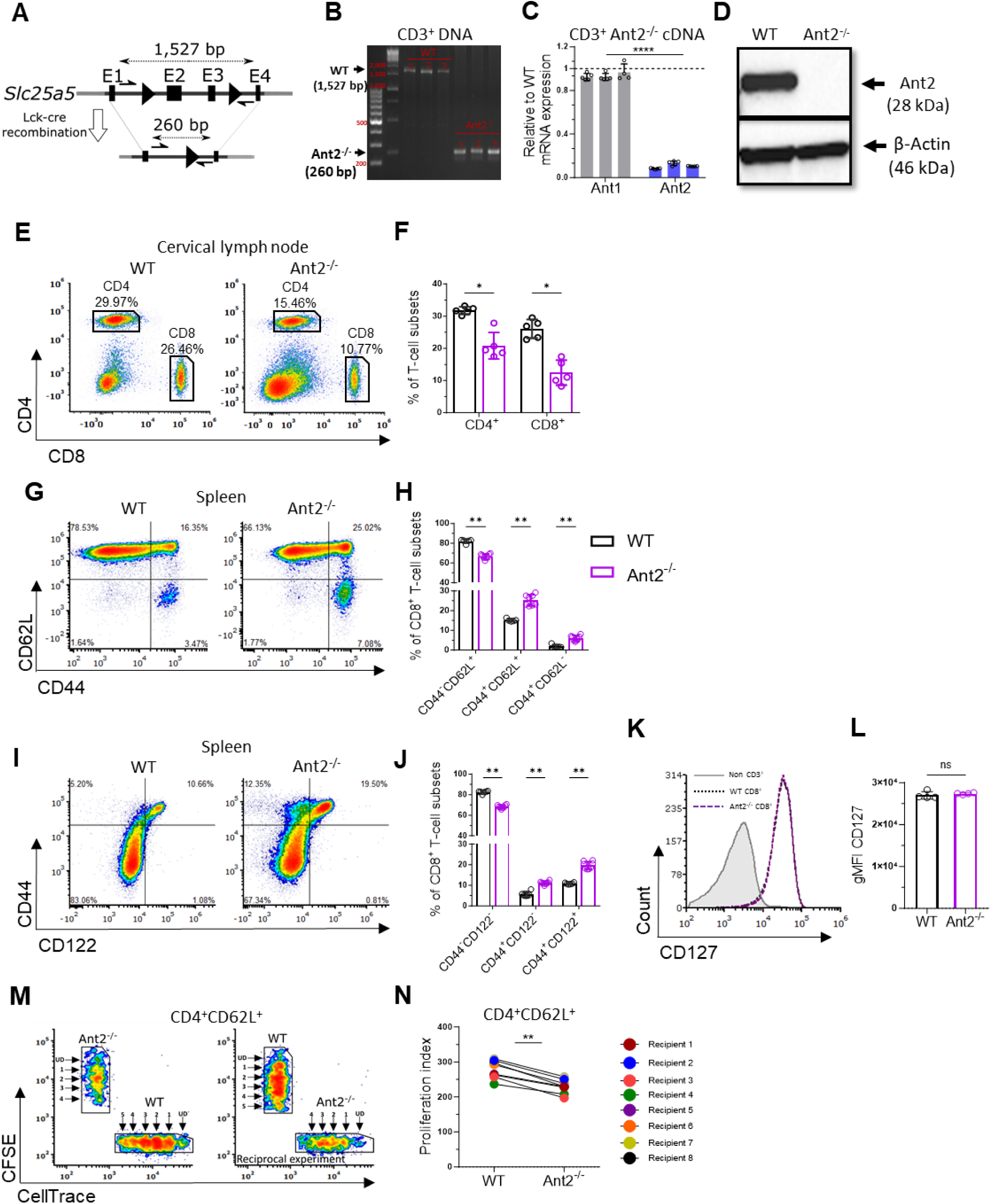
Lymphopenia in Ant2^−/−^ mice due to impaired homeostatic expansion of T cells. **(A-D)** Verification of T cell-specific deletion of the *Slc25a5* gene, encoding for Ant2 protein. **(A)** A schematic representation of *Slc25a5* nullification *in-vivo* using the cre-loxP system. Black triangle represents LoxP sequence, E=Exon. **(B)** PCR amplification of DNA confirming *Slc25a5* deletion in genomic DNA of purified CD3^+^ cells from Ant2^−/−^ and WT mice. Primers were designed to amplify intron 1 through intron 3, resulting in a 1,527 bp product for the WT “untouched” allele, and a 260 bp product for the successfully deleted exons 2+3. **(C)** Normalized mRNA expression of genes encoding for *Ant1* and *Ant2* in purified CD3^+^ Ant2^−/−^ cells relative to CD3^+^ WT cells as determined by real-time PCR. Normalization was performed against β2m expression. **(D)** Representative immunoblot analysis of whole-protein extracts from purified WT and Ant2^−/−^ CD3^+^ cells using an anti-Ant2 antibody, with an anti-β-actin antibody used as a loading control. **(E)** Representative flow cytometry plots of CD8 vs. CD4 in the cervical lymph node derived from 6-9 weeks old WT and Ant2^−/−^ littermate mice. Numbers indicate the frequencies of CD8^+^ and CD4^+^ T cells. **(F)** Bar graph summarizing the results shown in E. **(G)** Representative flow cytometry plots showing CD44 vs. CD62L gated on CD8^+^ T cells. **(H)** Bar graph summarizing the results shown in G. **(I)** Representative flow cytometry plots showing CD122 vs. CD44 gated on CD8^+^ T cells. **(J)** Bar graph summarizing the results shown in I. **(K)** Representative flow cytometry overlay histogram of CD127 intensity in WT and Ant2-/-CD8^+^ T cells. **(L)** Bar graph summarizing the results shown in K, as geometric mean fluorescence intensity (gMFI). **(M)** Representative flow cytometry plots of CellTrace vs CFSE intensities gated on donor CD4^+^ CD62L^hi^ T cells. **(N)** Graph depicting paired proliferation indexes of donor WT and Ant2^−/−^ CD4^+^ T cells. Statistical method: Two-tailed Mann–Whitney test (C, F, H, J, and L), or two-tailed Wilcoxon matched-pairs signed rank test (N). Data are represented as mean ± S.D. (P value, *P≤0.05, **P≤0.01, ****P≤0.0001).

**Supplementary Figure 2:**
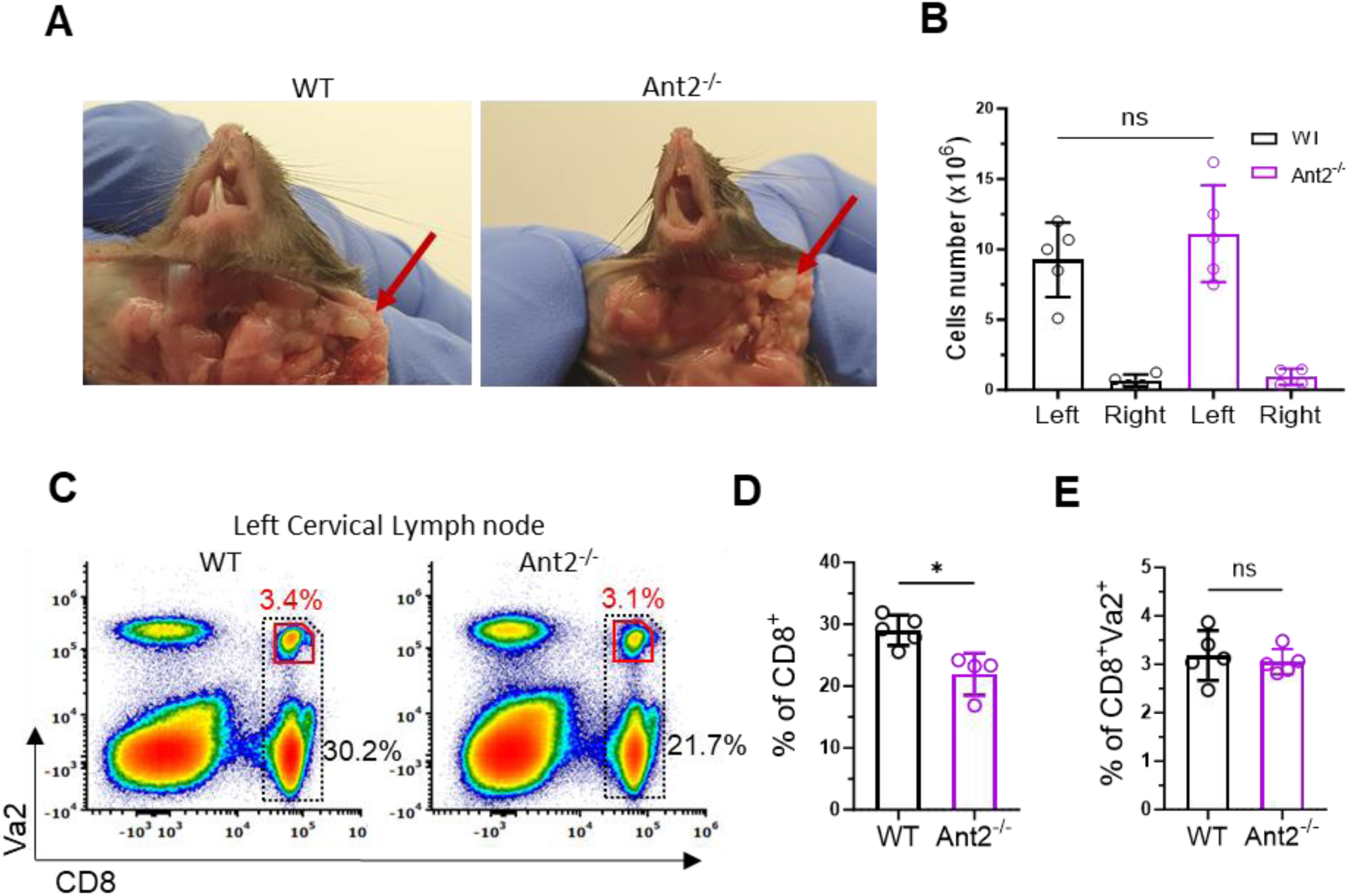
Intact T cell-mediated immunity in Ant2^−/−^ mice. **(A-E)** WT and Ant2^−/−^ littermate mice were primed intradermally in the left ear-pinna with 5 × 10^6^ transduction units (TU) of lentivirus expressing OVA (LvOVA). **(A)** Representative photos demonstrating the swelling of the left cervical lymph node (LN) compared to the right LN seven days post-infection (The uninfected right cervical LN served as a control for comparison). **(B)** Bar graphs show the absolute cell numbers of left and right cervical LNs. **(C)** Representative flow cytometry plots showing CD8 vs Vα2 staining from the cervical LNs of WT and Ant2^−/−^ mice. Numbers indicate the percentages of CD8^+^Vα2^+^ T cells (red font), and total CD8^+^ T cells (black font) **(D-E)** Bar graphs summarizing the results in C. Statistical method: Two-tailed Mann–Whitney test. Data are represented as mean ± S.D. (P value, *P≤0.05).

**Supplementary Figure 3:**
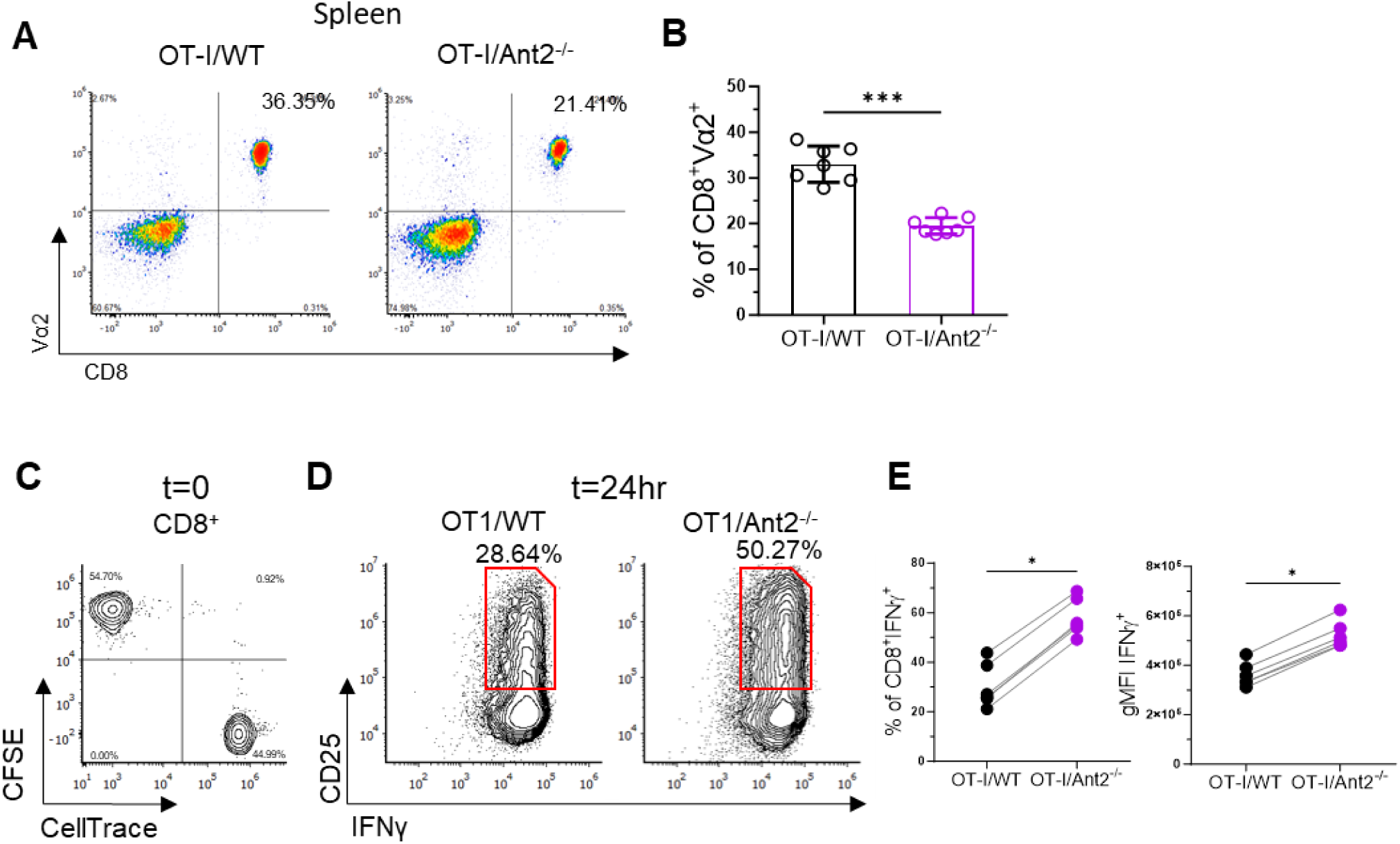
Increased activation potential of Ant2^−/−^ CD8^+^ T cells. **(A)** Representative flow cytometry plots showing CD8 vs Vα2 staining in spleens of WT and Ant2^−/−^ OT-I mice. **(B)** Bar graphs summarizing the results shown in A, as the percentage of CD8^+^ Vα2^+^ cells. **(C-E)** CFSE-labelled OT-I/WT and CellTrace-labelled OT-I/Ant2^−/−^ CD8^+^ T cells were co-cultured at approximately a 1:1 ratio and activated with 1µg/ml OVA-peptide. Twenty-four hours later, cells were subjected to intracellular staining for IFNγ. **(C)** Flow cytometry plot of CellTrace vs CFSE gated on CD8^+^ T cells at t=0. **(D)** Representative flow cytometry plots showing the frequency of IFNγ^+^CD25^+^ gated on CD8^+^ T cells. **(E)** The graphs summarize the results shown in D, as the frequency of CD8^+^IFNγ^+^ T cells **(left)**, and the gMFI of IFNγ gated on CD8^+^IFNγ^+^CD25^+^ T cells **(right)**. Each line represents both OT-I/WT (black dot) and OT-I/Ant2^−/−^ (purple dot) in the same well. Statistical method: Two-tailed Mann–Whitney test (B), or two-tailed Wilcoxon matched-pairs signed rank test (E). Data are represented as mean ± S.D. (P value, *P≤0.05, ***P≤0.001).

**Supplementary Figure 4:**
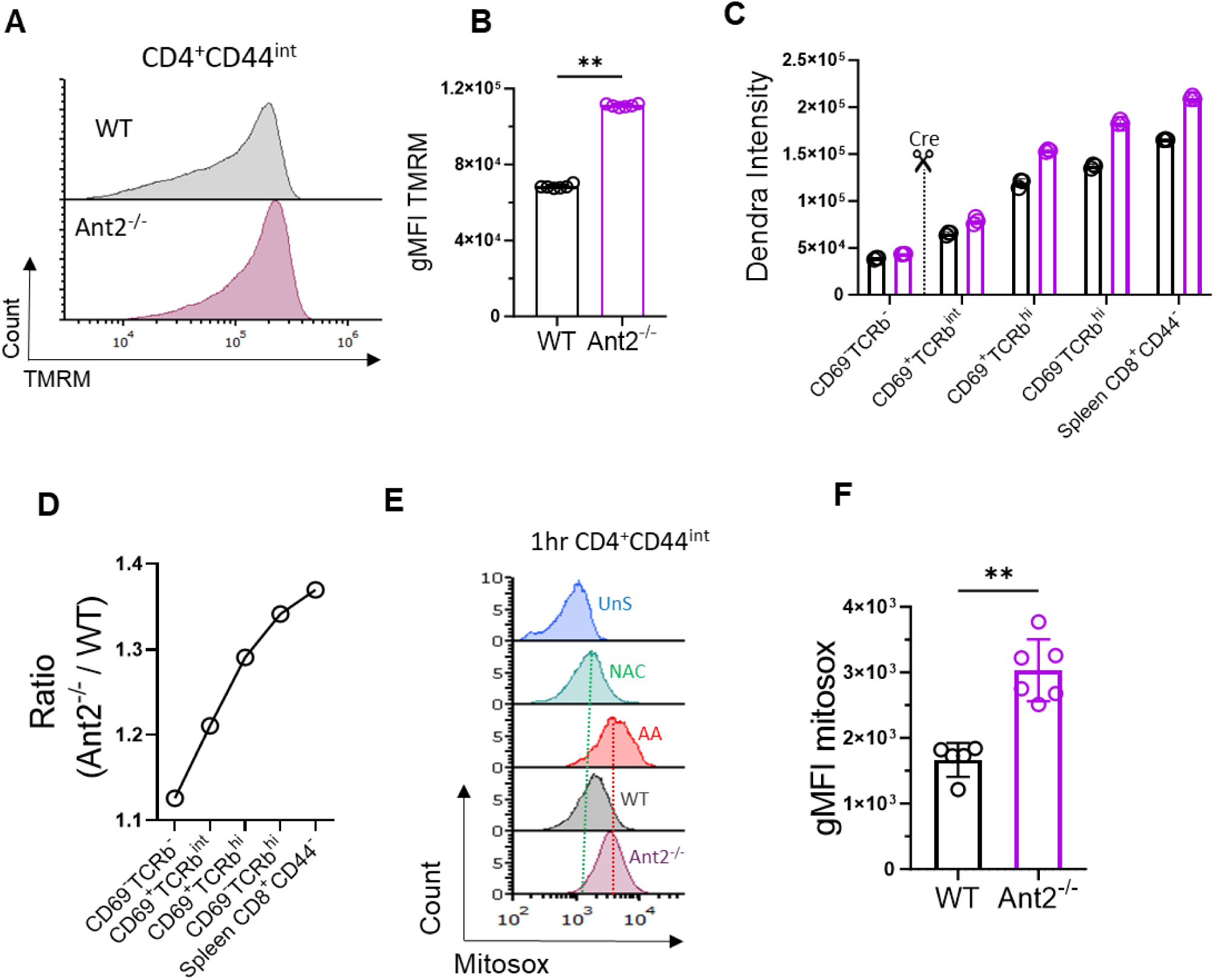
Increased mitochondrial biogenesis, membrane potential and ROS in Ant2^−/−^ thymocytes and CD4^+^ T cells. **(A)** Representative flow cytometry histogram of Tetramethylrhodamine, Methyl Ester, Perchlorate (TMRM) staining gated on naïve (CD44^int^) CD4^+^ T cells. **(B)** Bar graph summarizing the results shown in A. **(C-D)** Thymus and spleen from WT and Ant2^−/−^ mito-Dendra2 were harvested and analyzed using flow cytometry. Thymocytes were stained for CD69 and TCRβ. Splenocytes were stained for CD8 and CD44. **(C)** Bar graph summarizing the Dendra2 mean fluorescence intensity (gMFI) measured by flow cytometry in different stages of thymocyte development according to their CD69 and TCRβ expression (Stages A-D), as well as in the naïve (CD44^−^) CD8^+^ T cells in the spleen. **(D)** Graph representing the average ratio of Dendra2 intensity in Ant2^−/−^ CD8^+^ T cells compared to the corresponding stage in WT mice. This ratio is calculated by dividing the Dendra2 gMFI of Ant2^−/−^ cells in each stage by the Dendra2 gMFI of WT cells in the corresponding stage. **(E)** Representative flow cytometry histograms of MitoSOX® staining gated on one-hour stimulated CD4^+^(CD44^int^) T cells. WT CD4^+^ T cells were used as controls, with the following conditions from top to bottom: unstained (UnS), pre-treated with 200µM N-Acetyl Cysteine (NAC), and treated with 1µM Antimycin A (AA). **(F)** Bar graph summarizing the results shown in E as gMFI of MitoSOX®. Statistical method: Two-tailed Mann–Whitney test. Data are represented as mean ± S.D. (P value, **P≤0.01).

**Supplementary Figure 5:**
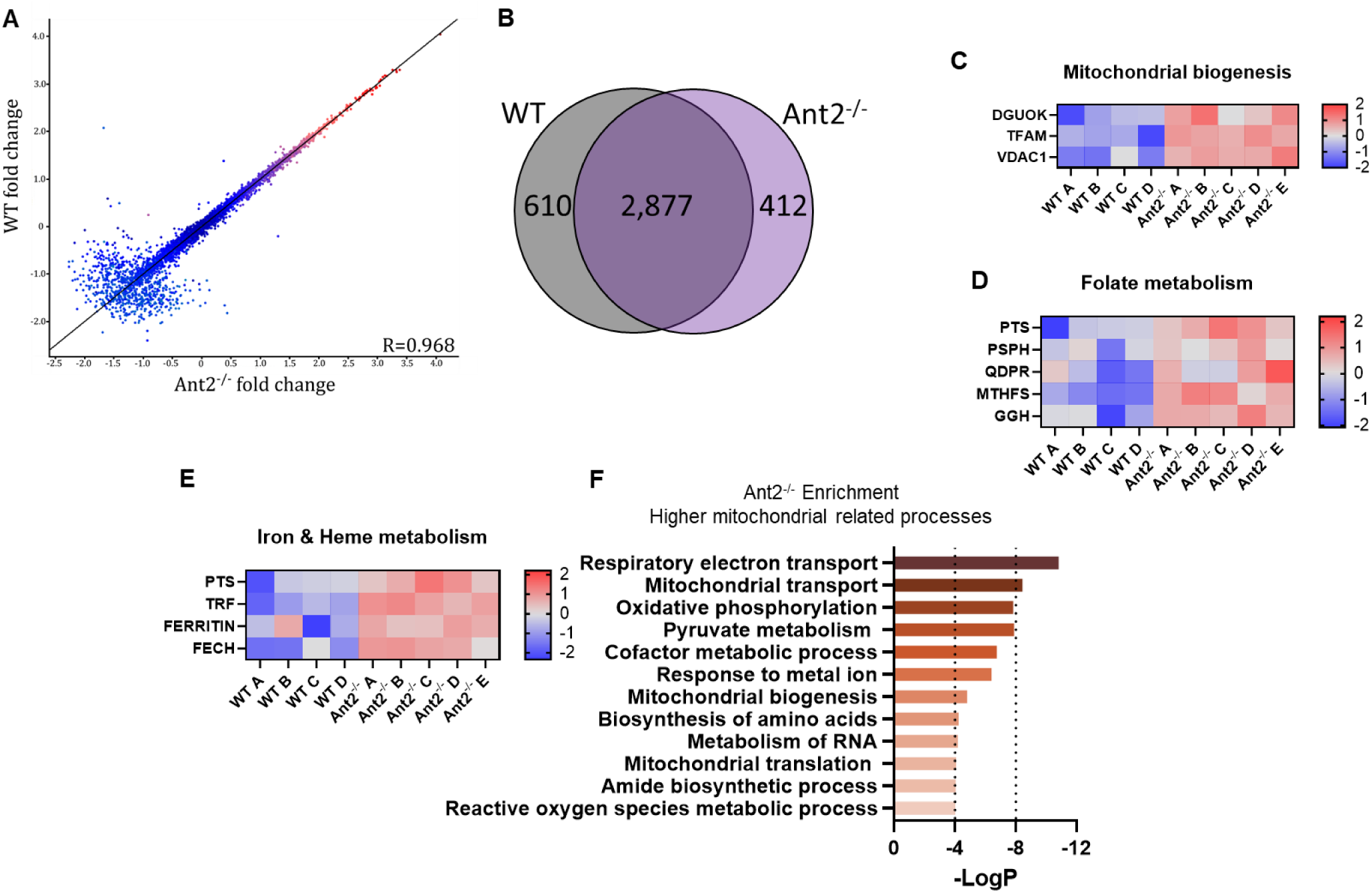
Altered proteome and distinct mitochondria in naïve Ant2^−/−^ CD8^+^ T cells. Mass-spectrometry analysis was performed on protein extracts from naïve (CD44^−^) CD8^+^ T cells derived from WT or Ant2^−/−^ OT-I mice (n=4 or 5 in each group). **(A)** Scatter plot of the Z-score normalized log_2_-transformed LFQ values of all 3,899 detected proteins in WT and Ant2^−/−^ naïve CD8^+^ T cells (p < 0.05). The Pearson correlation coefficient was calculated between the expression of each protein in the two samples. **(B)** Venn diagram plot showing relationships between all detected proteins in naïve WT and Ant2^−/−^ CD8^+^ T cells (p-value<0.05). **(C-E)** Heatmaps illustrating differentially expressed proteins associated with mitochondrial biogenesis **(C)**, Folate metabolism **(D)**, and iron & heme metabolism **(E)**. The fold change was determined through a student’s t-test analysis conducted on log_2_-transformed data after the Z-score normalization step. The color gradient spans from red, indicating a substantial fold change, to blue, representing a minimal fold change. **(F)** Summary of the pathway-enrichment analysis of all elevated proteins in Ant2^−/−^ adapted from Metascape.

**Supplementary Figure 6:**
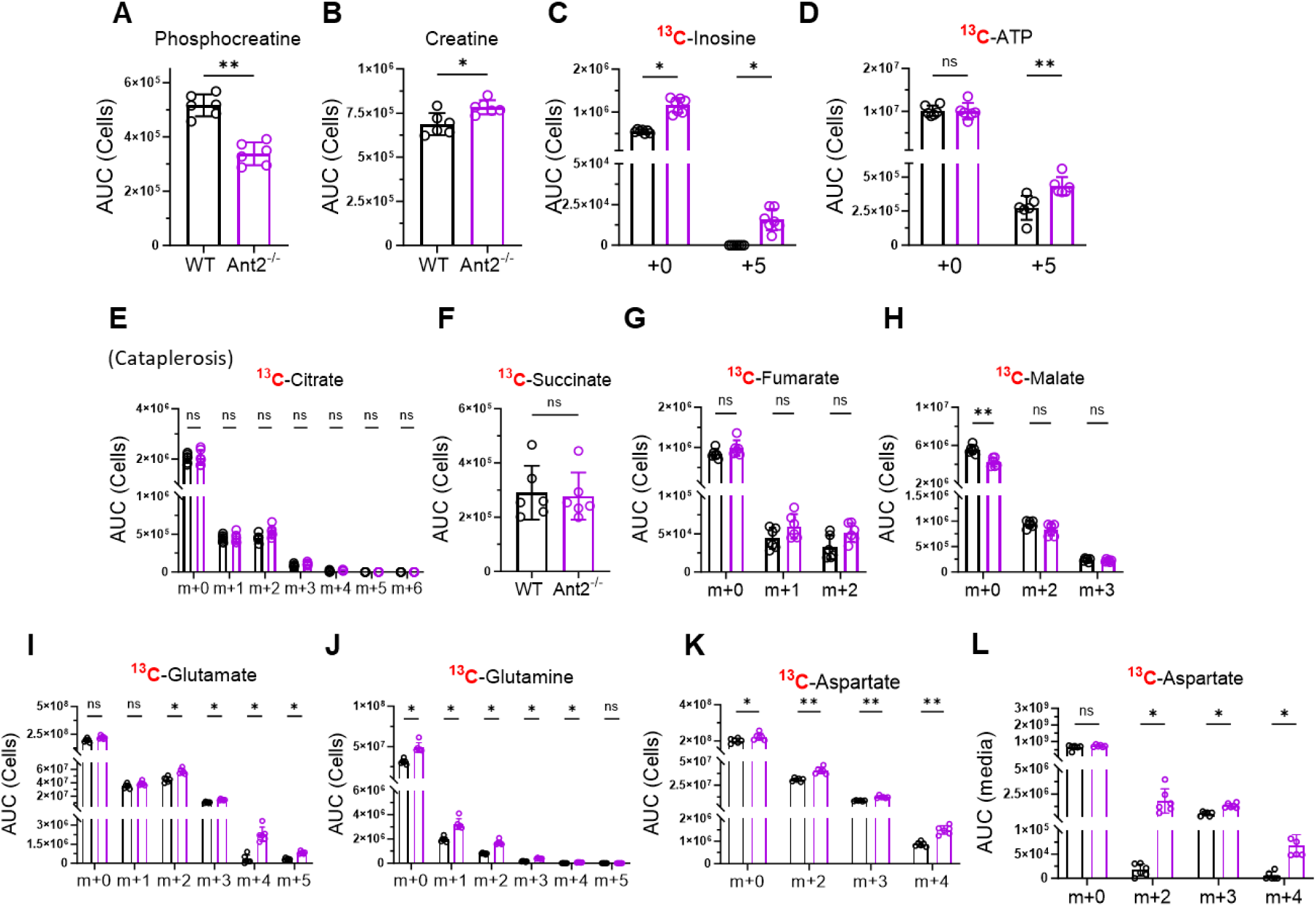
Metabolomic adaptations of naïve Ant2^−/−^ CD8^+^ T cells. **(A-M)** Comparison of the area under the curve (AUC) values obtained from metabolomics analysis combined with ^13^C_6_-glucose tracing between naïve (CD44^−^) WT and Ant2^−/−^ CD8^+^ T cells. **(A-B)** Bar graphs showing the AUC values of unlabeled phosphocreatine **(A)** and creatine **(B)** in the cell fraction. **(C-D)** Bar graph showing the AUC values of, unlabeled and m+5 inosine **(C)** and ATP **(D)** in the cell fraction. **(G-J)** Bar graphs displaying the AUC values of TCA cycle intermediates, citrate **(E)**, succinate **(F)**, fumarate **(G)**, and malate **(H)** labeling patterns in the cell fraction. **(G-J)** Bar graphs displaying the AUC values of TCA cycle derived amino-acids: glutamate **(I)**, glutamine **(J)**, and aspartate **(K)** labeling patterns in the cell fraction. Bar graphs showing the AUC values of aspartate in the media fraction **(L)**. Statistical method: Two-tailed Mann–Whitney test. Data are represented as mean ± S.D. (P value, *P≤0.05, **P≤0.01).

**Supplementary Figure 7:**
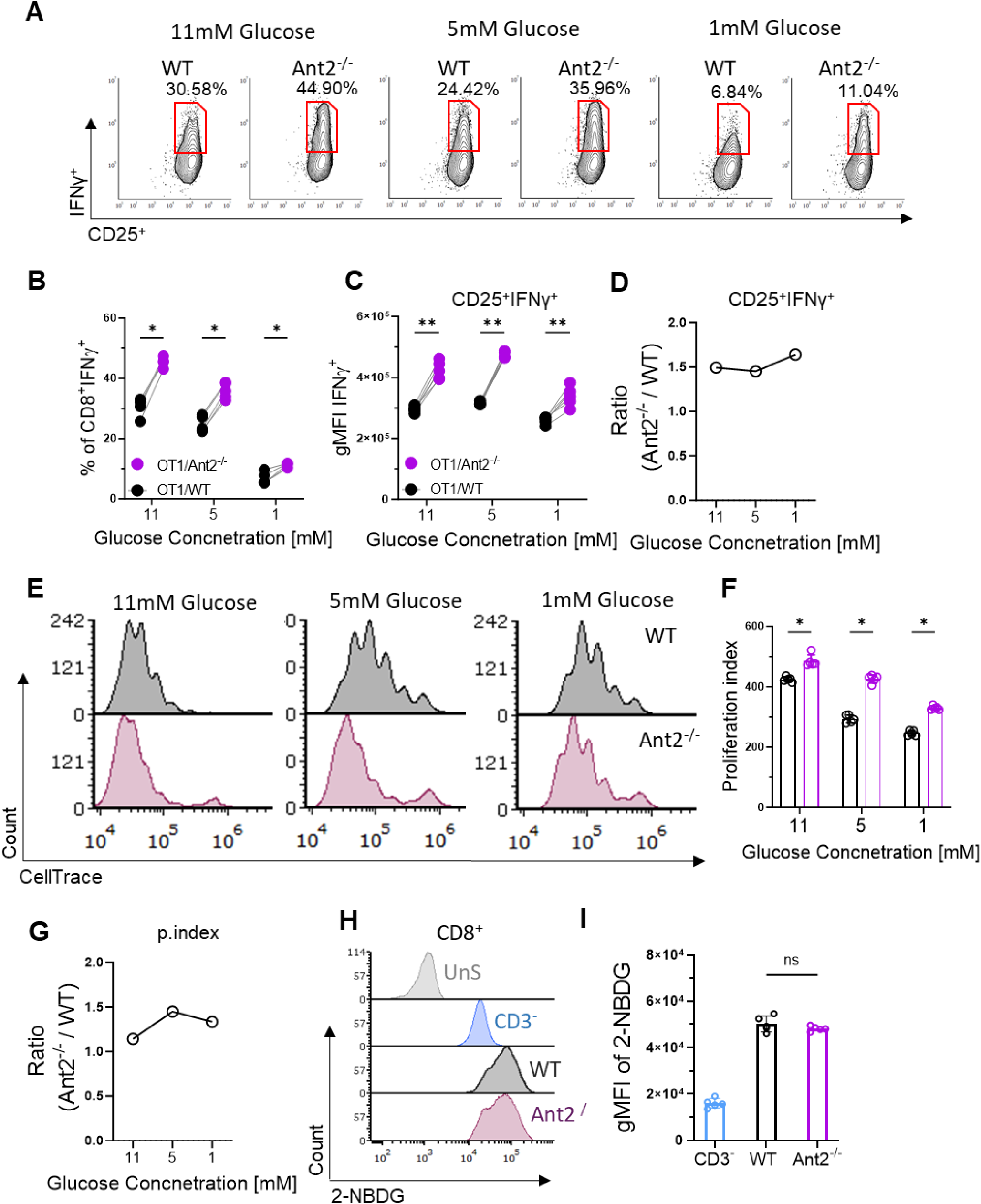
Metabolic alterations in Ant2-deficient T cells are not dependent on increased glucose concentration or uptake. **(A-B)** WT and Ant2^−/−^-derived splenocytes were CFSE- and CellTrace-labelled, respectively. CD8^+^ T cells were co-cultured in a ∼1:1 ratio using media supplemented with 11mM (control), 5mM, and 1mM glucose. Cells were activated with 100ng/ml αCD3 and 50ng/ml αCD28 soluble antibodies for 24h. **(A)** Representative flow cytometry plots of CD25 vs IFNγ gated on CD8^+^ T cells in media supplemented with reduced glucose concentrations as indicated. **(B-D)** Graphs summarizing the results in A, as depicted paired of the % of CD8^+^IFNγ^+^ **(B)**, and gMFI of IFNγ, gated on CD8^+^IFNγ^+^CD25^+^ **(C)**. Each line represents both WT (black dot) and Ant2^−/−^ (purple dot) CD8^+^ T cells co-cultured at the same well. **(D)** Graph representing the ratio of the average percentage of IFNγ^+^ CD8^+^ T cells derived from Ant2^−/−^ mice to WT mice for each corresponding glucose concentration. **(E-G)** CellTrace-labelled WT and Ant2^−/−^ derived splenocytes were cultured in media supplemented with 11mM (control), 5mM, and 1mM glucose. The cells were then activated as indicated above for 72h. **(E)** Representative flow cytometry histograms of CellTrace intensity gated on CD8^+^ T cells. **(F)** Bar graph summarizing the results of C, as proliferation index of WT and Ant2^−/−^ CD8^+^ T cells. **(G)** Graph representing the ratio of the average proliferation index of CD8^+^ T cells derived from Ant2^−/−^ mice to WT mice for each corresponding glucose concentration. **(H-I)** WT and Ant2^−/−^derived splenocytes were cultured in a glucose-free medium containing 60µM 2NDBG-FITC for 20 minutes. Cells were then analysed by flow cytometry. **(H)** Representative flow cytometry histograms of 2NDBG-FITC intensity gated on WT CD3^−^ (control), and WT and Ant2^−/−^ CD8^+^ T cells. **(I)** Bar graph summarizing the results of E, as gMFI of 2NDBG-FITC. Statistical method: Two-tailed Wilcoxon matched-pairs signed rank test (B-D), or by two-tailed Mann–Whitney test (F-H). Data are represented as mean ± S.D. (P value, *P≤0.05, **P≤0.01).

## Material and Methods

This study did not generate new unique reagents.

### Mice

The C57BL/6J (wild-type) mice (strain 000664), Slc25a5^tm1.1Nte^/J mice (Ant2) (strain 029482), B6.Cg-Tg(dLck-cre)3779Nik/J mice (strain 012837), C57BL/6-Tg(TcraTcrb)1100Mjb/J (OT-I) mice (strain 00383), and B6.SJL-*Ptprc*^a^ *Pepc*^b^/BoyJ mice (CD45.1) (strain 002014) were obtained from The Jackson Laboratory. The T cell-specific Ant2 knockout mice were generated by crossing mice containing a conditional floxed allele of Ant2 Slc25a5^tm1.1Nte^/J (Ant2^2f^) with transgenic mice expressing Cre under the control of the distal Lck gene promoter (dLck-Cre). B6;129S-Gt(ROSA)26Sor^tm1(CAG-COX8A/Dendra2)Dcc^/J (mito-Dendra2) mice (strain 018385) were a kind gift from Dr. Tsvee Lapidot from the Weizmann Institute of Science. The C57BL/6JOlaHsd mice used as recipient in the B16-OVA experiments were obtained from Envigo. All mice used for experiments at 6–14 weeks of age. Mice were maintained and bred under specific pathogen-free (SPF) conditions in the Hebrew University animal facilities according to Institutional Animal Care and Use Committee regulations.

### Antibodies

The following antibodies were used for flow cytometry: anti-CD8α (53–6.7), anti-CD4 (GK1.5), anti-CD3ε (145-2C11), anti-CD122 (5H4), anti-CD127 (A7R34), anti-CD62L (MEL-14), anti-CD44 (IM7), anti-CD69 (H1.2F3), anti-TCRβ (H57-597), anti-CD25 (PC61), anti-CD45.1 (A20), anti-CD45.2 (104), anti-TCR Vα2 (B20.1), anti-CD25, and anti-IFNγ (XMG1.2). All antibodies were purchased from BioLegend.

For the activation of mouse T cells, purified anti-CD3ε (145–2C11) and anti-CD28 (37.51) antibodies obtained from BioLegend were employed at the appropriate concentrations as per the experiment design. For immunoblotting, Antibodies to PDH phosphorylated on Ser^300^ (AP1064) and anti-PDH (ABS2082) were purchased from Sigma-Aldrich. The anti-Ant2 was acquired from Cell Signaling (14671). The anti-β-actin (sc-47778) and anti-β-tubulin (sc-5274) antibodies were purchased from Santa Cruz. These antibodies were diluted by 1:1,000 in TBST, with 3% BSA and 0.05% sodium azide.

### Polymerase Chain Reaction (PCR) for Ant2^−/−^ validation

Total DNA extraction was performed on CD3^+^ cells that were isolated from WT or Ant2^−/−^ mice using the ‘T cell isolation kit’ (StemCell^TM^, 19851). Cells were then lysed in a ‘Tail Buffer’ composed of 20 mM NaOH and 100µM EDTA (BI, 01-862-1B). The mixture was incubated at 95°C for 20 minutes. The PCR reaction was carried out by KAPA2G Fast ReadyMix™ (Sigma Aldrich, KK5103). Primers were designed targeting the intron 1 and intron 3 regions of the target DNA. The annealing temperature for the PCR was set to 65°C for 34 cycles.

F’ primer: ATGGTGCTGCTCAATTCTTAAACA,

R’ primer: AGCACAGGCATTGACTGGAGAACA.

### Quantitative real-time PCR and cDNA preparation

Total RNA was extracted from purified CD3^+^ cells that were isolated from WT or Ant2^−/−^ mice using the ‘T cell isolation kit’ (StemCell^TM^, 19851). Cells were then lysed using the Direct-zol™ RNA Mini Prep kit (Zymo Research). cDNA was synthesized from the isolated RNA using the ProtoScript® First Strand cDNA Synthesis Kit (Bio-Labs). Quantitative real-time PCR using Applied Biosystems (AB), Viia 7 Real-Time PCR system with a Power SYBR green PCR master mix kit (Applied Biosystems, 4367660) according to the manufacturer’s instructions.

The following primers were used to amplify the target genes:

Ant1 F’: GATCGAGAGGGTCAAACTGC, and R’: CTTTGAAGGCGAAGTTCAGG.

Ant2 F’: CCTTCGCCAAGGACTTCT, and R’: ATGACATTGGCCAGGTTCC.

### Mitochondrial DNA quantification

Total DNA (mitochondrial DNA and nuclear DNA) was extracted from CD8^+^ T cells that were isolated from WT or Ant2^−/−^ mice using the ‘CD8^+^ T cell isolation kit’ (StemCell^TM^, 19853). Cells were then lysed using the DNA extraction method (PCIA). Briefly, cell lysates were treated with lysis buffer composed of 10 mM Tris-HCl (pH 8), 1 mM EDTA, 0.5% SDS, and Proteinase K (80 units/ml) for overnight 55°C incubation. The lysates were then centrifuged, and the supernatant was treated with PICA buffer (Sigma Aldrich, P2069). The upper layer containing DNA was then carefully transferred to another Eppendorf tube and mixed with an equal volume of chloroform. After homogenization, the samples were centrifuged again, and the upper layer was transferred to a new tube. 3M NaAc was added (1:10), followed by an equivalent volume of cold (−20°C) isopropanol. The mixture was incubated at −20°C for 10 minutes and then centrifuged at 17,000 g for 10 minutes. The supernatant was discarded, and the DNA pellet was washed with 1 ml of 70% ethanol. After centrifugation, the residual ethanol was removed, and the DNA pellet was dissolved in 40 µl of pre-heated (55°C) double-distilled water (DDW). A concentration of 200 pg/ml of extracted DNA (total 800 pg/12 µl) was used in the RT-qPCR experiment.

β2M F’ primer: TGAGGCTTATTGCAATGCTG,

β2M R’ primer: ATGGCGGTTACAGTCCAAAG.

Mito-RNR2 F’ primer: CTAGAAACCCCGAAACCAAA,

Mito-RNR2 R’ primer: CCAGCTATCACCAAGCTCGT.

co-1 F’ primer: GCAGGAGCATCAGTAGACCTAAC,

co-1 R’ primer: GGAGTTTGATACTGTGTTATGGCTGG.

Data was normalized to Mouse endogenous control (β2M) and analyzed using ΔΔCt model. Each experiment was performed in eight replicates and was repeated three times.

### Western Blot

For anti-PDH immunoblot analysis, CD8^+^ T cells obtained from both WT and Ant2^−/−^ mice were isolated using a ‘CD8^+^ T cell isolation kit’ (StemCell^TM^, 19853). Alternatively, CD3^+^ cells were isolated using a ‘T cell isolation kit’ (StemCell^TM^, 19851) for anti-Ant2 immunoblot. Subsequently, the isolated cells were lysed in RIPA buffer (Thermo Fisher Scientific, 89900) supplemented with protease and phosphatase inhibitors (Thermo Fisher Scientific, 78440). The quantification of protein content was conducted using the Bradford assay (Bio-Rad, 5000001). A total of 30 µg of protein was subjected to separation by SDS-PAGE and transferred to nitrocellulose membranes. Membranes were blocked with 5% BSA in TBST for 1 hour at room temperature. After blocking, membranes were washed three times with TBST and incubated with primary antibodies overnight at 4°C. After washing, membranes were incubated with horseradish peroxidase-conjugated secondary antibodies (Abcam, Ab98799 or 97085) for 30 minutes.

### Flow cytometry

Cells were stained for cell-surface markers using various conjugated monoclonal antibodies (mAbs) in FACS buffer (PBS supplemented with 2% FBS and 1 mM EDTA) for 30 minutes at 4°C.

For the staining of mitochondrial membrane potential, cells were treated with TMRM (Molecular Probes, Eugene, OR) at a concentration of 50 nM in FACS buffer without EDTA for 30 minutes at 37°C. Following this, the samples were maintained on ice for surface staining until they were ready for analysis. For intracellular staining, cells were fixed in 1% paraformaldehyde (PFA) in PBS for 15 minutes at RT. Cells were then washed with PBS, and permeabilized using a solution containing 0.1% saponin and intra-cellular mAbs in cold FACS buffer for 30 minutes.

Stained cells were analyzed by a CytoFLEX Flow Cytometer with data being captured and processed using the CytExpert Software (Beckman Coulter, Inc). Subsequent analysis of the collected data was performed using FACS Express 6 software (De Novo Software).

### *In-vivo* T cell homeostatic expansion assay

CD45.1^+/+^ WT recipient mice were sub-lethally irradiated (600 rad). One day following irradiation, an intravenous co-adoptive transfer was conducted. This involved injecting 4M/200µl T cells in a 1:1 ratio of CD45.2^+/+^ CFSE-labeled WT and CellTrace (Thermo Fischer Scientific, M20036) labeled-Ant2^−/−^-derived T cells, into the sub-lethally irradiated recipient mice. Reciprocal labeling was also performed. After 7 days, the mice were euthanized, and their splenocytes were isolated and examined by flow cytometry for CFSE and CellTrace intensities gated on naive (CD62L^hi^) CD4^+^ and CD8^+^ donor T cells.

### *In-vivo* immunization assay for testing CD4^+^ mediated immunity

WT and Ant2^−/−^ mice were subcutaneously immunized three times at 7-day intervals. The first two immunizations included 100µg/100µl OVA protein (Sigma Aldrich, 9006-59-1) in complete Freund’s adjuvant (CFA) (Sigma Aldrich, F5881), while the third immunization included OVA protein without CFA. After 21 days from the first vaccination, sera were collected and tested for OVA-specific IgG1 (Biolegend, 1144-05) & IgG2 (Biolegend, 1155-05) antibodies, as well as for the IFNγ (Biolegend, BLG-430804) and IL-4 (Biolegend, BLG-431104) cytokines using enzyme-linked immunosorbent assay (ELISA).

### *In-vivo* CD8^+^ T cell activation assay using lentiOVA

WT or Ant2^−/−^ mice were primed intradermally in the ear pinna with 5 × 10^6^ transfection units (TU) of Lv-OVA-GFP (A gift from Dr. Avihai Hovav from the Hebrew University of Jerusalem). Seven days after the viral challenge, mice were sacrificed, and their left and right cervical lymph nodes were dissected and analyzed by a flow cytometer.

### *In-vivo* T cell activation assay

CD45.1^+/+^ WT recipient mice were intravenously (i.v) administered CellTrace-labeled CD45.2^+/+^ 2M/200µl CD8^+^ T cells derived from WT or Ant2^−/−^ OT-1 mice. After one day, recipient mice were intraperitoneally (i.p) injected with 50µg of OVA protein (Sigma Aldrich, 9006-59-1). On day three following cell transfer, mice were euthanized, and their splenocytes were isolated and examined by flow cytometry for CellTrace intensity.

### *Testing effects of* CATR treatment on T cell activation *In-vivo*

Littermate OT-I mice were administered either carboxyatractyloside (CATR) at a dose of 2mg/kg or vehicle for a duration of 2 weeks. CD8^+^ T cells from the donor mice were labeled with Cell-Trace and subsequently intravenously administered to recipient WT mice. On the following day, the recipient mice were injected intraperitoneally with either 25µg or 100µg of OVA (Sigma Aldrich, 9006-59-1), as per the experimental design. After 3 days, the mice were euthanized, and their spleens were analyzed using flow cytometry to assess CellTrace intensity and IFNγ production. For IFNγ production, 1M/100µl splenocytes (flat 96-well plate) derived from OT-1 mice treated with either CATR or vehicle were restimulated with 1µg/ml OVA-peptide for 6 hours and Brefeldin A (Biolegend, 420601) for 4hr prior to analysis.

### *In-vivo* B16-OVA melanoma tumor model

Donor WT OT-I mice received daily IP injections of either PBS (mock) or 1 mg/kg ATR for 10 days. Concurrently, 6-week-old female CD45.2 C57BL/6 recipient mice (Envigo) were anesthetized with ketamine & xylazine diluted in PBS, shaved, and injected intradermally on the right flank with 1 x 10^6^ B16-OVA tumor cells in 100μL HBSS (n=4/group). At day 6 post-tumor injection, donor mice were sacrificed, and their T cells were activated in vitro with 0.5 µg/ml SIINFEKEL-peptide for 2 days. At day 7 post-tumor injection (pti), recipient mice were sub-lethally irradiated (550 rad). At day 8 pti, activated donor OT-1 T cells were adoptively transferred to the recipient mice. Tumor volumes were measured every 2 days. Mice were euthanized when tumors reached >1400 mm^3^ or at the study endpoint (day 26).

### *In-vitro* T cell for proliferation and cytokines production assay

Splenocytes derived from both WT and Ant2^−/−^ mice were labeled with CellTrace and incubated with appropriate amounts of suspended anti-CD3ε and half the amount of anti-CD28 antibodies, as per the experiment design. The incubation was in RPMI (Sigma Aldrich, 11875093) containing 10% Heat inactivated Fetal Bovine Serum (Gibco, 10500064), 2mM Glutamine (Satorius, 03-042-1B), 100 U/ml penicillin-streptomycin (Satorius, 03-031-1B), and 50µM β-mercaptoethanol (Merck, 60-24-2). For the proliferation assay, cells were incubated for 72 hours before analysis. The proliferation index reflects the sum of the percentage of cells in each generation group multiplied by the number of divisions. For the IFNγ assay, cells were activated for 24 hours and treated with Brefeldin A (Biolegend, 420601) 3 hours before analysis.

### *In-vitro* T cell killing assay

CD8^+^ T cells were negatively selected using StemCell^TM^ kit (19853) from WT and Ant2^−/−^-derived OT-I mice. Cells were activated in a 24-well plate coated with 2ug/ml of αCD3 and αCD28 antibodies for 5 days. Following this activation, the cells were incubated with either B16 (control) or B16 melanoma cells expressing ovalbumin (B16-OVA) target cells, maintaining a 1:1 ratio. After 6 hours, the cells underwent three PBS washes. The remaining cells were subjected to an MTT assay. The percentage of killing was determined using the formula: % of killing = 100 – {[(O.D. in B16-OVA with T cells -O.D. blank well) / O.D. in B16-OVA without T cells] x 100}. The calculated killing percentages were normalized to the control. Statistical analysis was conducted using the non-parametric Mann–Whitney test. The results are presented as mean ± SD, with P values indicated as *<0.01 and **<0.001.

### Seahorse assay

Extracellular flux experiments were performed using a Seahorse XFe96 Extracellular Flux Analyzer (Seahorse Bioscience). CD8^+^ T cells were negatively selected using StemCell^TM^ kit (19853) from WT and Ant2^−/−^-derived OT-I mice. Naïve or activated CD8^+^ T cells (4 × 10^5^ or 2 × 10^5^ cells per well, respectively) were seeded in Seahorse XF 96 designated plates using 100 µg/ml poly-D-lysine (Sigma-Aldrich, A-003-E) and assayed according to the manufacturer’s instructions. Briefly, RPMI 1640 medium (Bio-Rad, 11-100-1K, without sodium bicarbonate) was supplemented with 2 mM glutamine (Satorius, 03-042-1B), 1 mM sodium pyruvate (Satorius, 03-022-1B), and 100 U/ml penicillin-streptomycin (Satorius, 03-031-1B). The following compound concentrations were used for the OCR measurements: 2µM Oligomycin (Cayman Chemicals, 11342), 2µM FCCP (Cayman Chemicals, Cat# 15218), and 1µM Antimycin-A (Cayman Chemicals, 19433) & Rotenone (Cayman Chemicals, 13995). A total of 4 measurements per compound.

### MitoSOX**®** staining

WT and Ant2^−/−^ splenocytes were incubated with pre-heated 1.5µM MitoSOX red (abcam, ab228567) at 37°C for 20 minutes.

For control cells, a subset was pre-treated with 200µM NAC (negative control) or 1µM Antimycin A (positive control) in the incubator at 37°C for 10 minutes using Eppendorf tubes. Subsequently, cells were washed with pre-warmed (37°C) HBSS (from the same kit, ab228567) and immediately stained for MitoSOX^TM^ along with the rest of the samples in a 96U well plate. After incubation, cells were washed twice in pre-warmed (37°C) HBSS and then subjected to surface staining with FACS buffer using the indicated antibodies.

### Confocal microscopy

CD8^+^ T cells were negatively selected using StemCell^TM^ kit (19853) from WT and Ant2^−/−^-derived mice. Cells were then seeded in high resolution chamber (1.5H, 170µm ± 5µm) (Ibidi, IBD-80807) in FACS buffer. Pictures were obtained using a Zeiss LSM 980 with Airyscan on a Zeiss Axio Observer 7 SP inverted microscope using a Plan-Apochromate 63×/1.4 Oil DIC objective. Analysis was performed using the NIS-Elements (Nikon)

### Protein mass spectrometry

#### LC-MS analysis

Analysis was performed at the Stein Family mass spectrometry center in the Silberman Institute of Life Sciences, Hebrew University of Jerusalem.

#### Sample preparation

Naïve CD8^+^ T cells were negatively selected using StemCell^TM^ kit (19858) from both WT and Ant2^−/−^-derived OT-I mice. Cell lysates were sonicated in sonication bath for 15 minutes. Lysates were cleaned and digested using S-Trap microcolumns (Protifi, LLC, Huntington, NY) as specified by the manufacturer. In brief: DTT was added to samples to final concentration of 10mM and incubated for 30 minutes at 37°C. The proteins were alkylated in 55 mM iodoacetamide and incubated for 30 minutes at room temperature in the dark. Protein samples were acidified using phosphoric acid at a final concentration of 1.2%. Methanol-TRIS buffer (90% MeOH, 10% TRIS 0.5M pH7.1) was added to the samples at a ratio of 6:1 (buffer: sample) and loaded onto S-Trap columns and subsequently washed twice with buffer. Sequencing grade modified trypsin (Promega Corp., Madison, WS) was loaded onto the column (1µg per column) and incubated at 47°C for one hour. Peptides were eluted from column, acidified, and desalted on homemade C18 stage tips (Rappsilber J, Mann M, Ishihama Y. Protocol for micro-purification, enrichment, pre-fractionation and storage of peptides for proteomics using StageTips. Nat Protoc. 2007;2(8):1896-906.). A total of 0.75 µg of peptides (determined by Absorbance at 280 nm) from each sample were injected into the mass spectrometer.

#### LC-MS/MS analysis

MS analysis was performed using a Q Exactive-HF mass spectrometer (Thermo Fisher Scientific, Waltham, MA USA) coupled on-line to an Ultimate 3000 Dionex (Thermo Fisher Scientific, Waltham, MA USA) UHPLC. Peptides were separated on a 120 min acetonitrile gradient run at a flow rate of 0.15 μl/min on a reverse phase 25-cm-long C18 column (75 μm ID, 2 μm, 100Å, Thermo PepMapRSLC). Survey scans (300–1,650 m/z, target value 3E6 charges, maximum ion injection time 20 ms) were acquired and followed by higher energy collisional dissociation (HCD) based fragmentation (normalized collision energy 27). A resolution of 60,000 was used for survey scans and up to 15 dynamically chosen most abundant precursor ions, with “peptide preferable” profile were fragmented (isolation window 1.6 m/z). The MS/MS scans were acquired at a resolution of 15,000 (target value 1E5 charges, maximum ion injection times 25 ms). Dynamic exclusion was 20 sec. Data were acquired using Xcalibur software (Thermo Scientific). To avoid a carryover, the column was washed with 80% acetonitrile, 0.1% formic acid for 25 min between samples.

#### MS data analysis

Mass spectra data were processed using the MaxQuant computational platform, version 1.6.6.0. Peak lists were searched against Mus musculus proteome obtained from Uniprot on 18.12.2019. The search included cysteine carbamidomethylation as a fixed modification, N-terminal acetylation and oxidation of methionine as variable modifications and allowed up to two miscleavages. The ‘match-between-runs’ option was used. Peptides with a length of at least seven amino acids were considered and the required FDR was set to 1% at the peptide and protein level. Relative protein quantification in MaxQuant was performed using the label-free quantification (LFQ) algorithm (Cox, J. et al. MaxLFQ allows accurate proteome-wide label-free quantification by delayed normalization and maximal peptide ratio extraction. Mol. Cell. Proteomics 13, 2513–2526 (2014).

### Metabolomics and isotope tracing analysis

#### Sample preparation

Naïve CD8^+^ T cells were negatively selected using StemCell^TM^ kit (19858) from WT and Ant2^−/−^-derived OT-I mice. The cells were incubated with either 11mM ^13^C_6_-Glucose or 2mM ^13^C_5_-Glutamine for 4 hours. Media extract: 40µl of culture medium was added to 40 μl of a cold extraction solution (−20°C) composed of methanol, acetonitrile, and water (5:3:2). Cell extracts: Cells were rapidly washed three times with ice-cold PBS, after which intracellular metabolites were extracted with 100μl of ice-cold extraction solution for 5 min at 4°C and subjected to 3 freeze-thaw cycles.

Media and cell extracts were centrifuged (10 min at 17,000 g, 4°C) to remove insoluble material, and the supernatant was collected for LC-MS analysis. Metabolomics data was normalized to protein concentrations using Bradford assay.

#### LC-MS analysis

Briefly, the supernatants from the media and cell extracts were analyzed using a Thermo Ultimate 3000 high-performance liquid chromatography (HPLC) system coupled to a Q-Exactive Orbitrap Mass Spectrometer (Thermo Fisher Scientific). The resolution was 35,000 at 200 mass/charge ratio (m/z), and the electrospray ionization (ESI) mode was used with polarity switching to enable both positive and negative ions across a mass range of 67 to 1000 m/z.

#### HPLC setup

The HPLC setup consisted of a ZIC-pHILIC column (SeQuant; 150 mm x 2.1 mm, 5 µm; Merck), with a ZIC-pHILIC guard column (SeQuant; 20 mm x 2.1 mm). 5 µl of biological extracts were injected, and the compounds were separated with a mobile phase gradient of 15 minutes, starting at 20% aqueous (20 mM ammonium carbonate adjusted to pH 2 with 0.1% of 25% ammonium hydroxide) and 80% organic (acetonitrile) and terminated with 20% acetonitrile. The flow rate and column temperature were maintained at 0.2 ml/min and 45°C, respectively, for a total run time of 27 minutes.

### Data acquisition and analysis

All metabolites were detected using mass accuracy below five parts per million (ppm). Thermo Xcalibur was used for data acquisition. TraceFinder 4.1 was used for analysis. Peak areas of metabolites were determined by using the exact mass of the singly charged ions. The retention time of metabolites was predetermined on the pHILIC column by analyzing an in-house mass spectrometry metabolite library that was built by running commercially available standards.

### Data analysis and statistics

The statistical significance of differences was determined by the two-tailed Mann-Whitney non-parametric t-test or two-tailed Wilcoxon matched-pairs signed rank test. Biological replicates refer to independent experimental replicates sourced from different mice donors. Technical replicates refer to independent experimental replicates from the same biological source. Differences with a P value of less than 0.05 were considered statistically significant. Graph prism and Perseus programs were used. MS data was normalized by ranking. Non-values were plugged with imputation from a normal distribution to prevent zeros bias.

## Author contributions

O.Y. and M.B. designed and performed research, analyzed data and wrote the manuscript; L.C-D, O.S, Z.B, V.L, A.P, I.A, B.A. performed research. A.S, E.G, M.H, J.T. analyzed data.

## Acknowledgments

This work was supported by grants from the Israel Science Foundation No.883/22 and No.2377/20, and the ICRF–CRI Immunotherapy Project Grant, a medical research grant from Israel Cancer Research Fund and Cancer Research Institute.

## Declaration of interests

The authors declare no competing interests.

## Declaration of generative AI and AI-assisted technologies in the writing process

During the preparation of this work the authors used Chat-GPT3.5 in order to improve language, grammar and readability. After using this tool, the authors reviewed and edited the content as needed and take full responsibility for the content of the publication.

## Notes

### Competing Interest Statement

The authors have declared no competing interest.

### Summary of Updates

The title of the article changed to fit the new results better. -Abstract changed accordingly. -Major changes to the introduction for better clarity. -Minor changes to the discussion.

## References

1. Reina-Campos, M., Scharping, N.E., and Goldrath, A.W. (2021). CD8+ T cell metabolism in infection and cancer. Nature Reviews Immunology 2021 21:11 21, 718–738. 10.1038/s41577-021-00537-8.

2. Pearce, E.L., Poffenberger, M.C., Chang, C.-H., and Jones, R.G. (2013). Fueling Immunity: Insights into Metabolism and Lymphocyte Function. Science (1979) 342. 10.1126/science.1242454.

3. Chapman, N.M., Boothby, M.R., and Chi, H. (2020). Metabolic coordination of T cell quiescence and activation. Nat Rev Immunol 20, 55–70. 10.1038/s41577-019-0203-y.

4. Bantug, G.R., Galluzzi, L., Kroemer, G., and Hess, C. (2018). The spectrum of T cell metabolism in health and disease. Nat Rev Immunol 18, 19–34. 10.1038/nri.2017.99.

5. Wang, R., and Green, D.R. (2012). Metabolic checkpoints in activated T cells. Nat Immunol 13, 907–915. 10.1038/ni.2386.

6. Klein Geltink, R.I., Kyle, R.L., and Pearce, E.L. (2018). Unraveling the Complex Interplay Between T Cell Metabolism and Function. 10.1146/annurev-immunol.

7. McBride, H.M., Neuspiel, M., and Wasiak, S. (2006). Mitochondria: More Than Just a Powerhouse. Current Biology 16, R551–R560. 10.1016/j.cub.2006.06.054.

8. Martínez-Reyes, I., Diebold, L.P., Kong, H., Schieber, M., Huang, H., Hensley, C.T., Mehta, M.M., Wang, T., Santos, J.H., Woychik, R., et al. (2016). TCA Cycle and Mitochondrial Membrane Potential Are Necessary for Diverse Biological Functions. Mol Cell 61, 199–209. 10.1016/J.MOLCEL.2015.12.002.

9. Ron-Harel, N., Santos, D., Ghergurovich, J.M., Sage, P.T., Reddy, A., Lovitch, S.B., Dephoure, N., Satterstrom, F.K., Sheffer, M., Spinelli, J.B., et al. (2016). Mitochondrial Biogenesis and Proteome Remodeling Promote One-Carbon Metabolism for T Cell Activation. Cell Metab 24, 104–117. 10.1016/j.cmet.2016.06.007.

10. Luengo, A., Li, Z., Gui, D.Y., Sullivan, L.B., Zagorulya, M., Do, B.T., Ferreira, R., Naamati, A., Ali, A., Lewis, C.A., et al. (2021). Increased demand for NAD+ relative to ATP drives aerobic glycolysis. Mol Cell 81, 691. 10.1016/J.MOLCEL.2020.12.012.

11. Saragovi, A., Abramovich, I., Omar, I., Arbib, E., Toker, O., Eyal, G., and Berger, M. (2020). Systemic hypoxia inhibits T cell response by limiting mitobiogenesis via matrix substrate-level phosphorylation arrest. Elife 9, 1–50. 10.7554/eLife.56612.

12. Gropper, Y., Feferman, T., Shalit, T., Salame, T.-M., Porat, Z., and Shakhar, G. (2017). Culturing CTLs under Hypoxic Conditions Enhances Their Cytolysis and Improves Their Anti-tumor Function. Cell Rep 20, 2547– 2555. 10.1016/j.celrep.2017.08.071.

13. Veliça, P., Cunha, P.P., Vojnovic, N., Foskolou, I.P., Bargiela, D., Gojkovic, M., Rundqvist, H., and Johnson, R.S. (2021). Modified Hypoxia-Inducible Factor Expression in CD8+ T Cells Increases Antitumor Efficacy. Cancer Immunol Res 9, 401–414. 10.1158/2326-6066.CIR-20-0561.

14. Cho, J., Seo, J., Lim, C.H., Yang, L., Shiratsuchi, T., Lee, M.H., Chowdhury, R.R., Kasahara, H., Kim, J.S., Oh, S.P., et al. (2015). Mitochondrial ATP transporter Ant2 depletion impairs erythropoiesis and B lymphopoiesis. Cell Death Differ 22, 1437–1450. 10.1038/cdd.2014.230.

15. Kunji, E.R.S., Aleksandrova, A., King, M.S., Majd, H., Ashton, V.L., Cerson, E., Springett, R., Kibalchenko, M., Tavoulari, S., Crichton, P.G., et al. (2016). The transport mechanism of the mitochondrial ADP/ATP carrier. Biochim Biophys Acta Mol Cell Res 1863, 2379–2393. 10.1016/j.bbamcr.2016.03.015.

16. Kieper, W.C., and Jameson, S.C. (1999). Homeostatic expansion and phenotypic conversion of naïve T cells in response to self peptide/MHC ligands. Proceedings of the National Academy of Sciences 96, 13306–13311. 10.1073/pnas.96.23.13306.

17. Surh, C.D., and Sprent, J. (2000). Homeostatic T Cell Proliferation. J Exp Med 192, F9–F14. 10.1084/jem.192.4.F9.

18. Bedard, M., and Shah, D.K. T-cell development in the thymus.

19. Bronstein-Sitton, N., Cohen-Daniel, L., Vaknin, I., Ezernitchi, A. V, Leshem, B., Halabi, A., Houri-Hadad, Y., Greenbaum, E., Zakay-Rones, Z., Shapira, L., et al. (2003). Sustained exposure to bacterial antigen induces interferon-γ-dependent T cell receptor ζ down-regulation and impaired T cell function. Nat Immunol 4, 957–964. 10.1038/ni975.

20. Furmanov, K., Elnekave, M., Sa’eed, A., Segev, H., Eli-Berchoer, L., Kotton, D.N., Bachrach, G., and Hovav, A.-H. (2013). Diminished Memory T-Cell Expansion Due to Delayed Kinetics of Antigen Expression by Lentivectors. PLoS One 8, e66488. 10.1371/journal.pone.0066488.

21. Clarke, S.Rm.K., Barnden, M., Kurts, C., Carbone, F.R., Miller, J.F., and Heath, W.R. (2000). Characterization of the ovalbumin-specific TCR transgenic line OT-I: MHC elements for positive and negative selection. Immunol Cell Biol 78, 110–117. 10.1046/j.1440-1711.2000.00889.x.

22. Budhu, S., Loike, J.D., Pandolfi, A., Han, S., Catalano, G., Constantinescu, A., Clynes, R., and Silverstein, S.C. (2010). CD8+ T cell concentration determines their efficiency in killing cognate antigen–expressing syngeneic mammalian cells in vitro and in mouse tissues. Journal of Experimental Medicine 207, 223–235. 10.1084/jem.20091279.

23. Desdín-Micó, G., Soto-Heredero, G., and Mittelbrunn, M. (2018). Mitochondrial activity in T cells. Preprint at Elsevier B.V., 10.1016/j.mito.2017.10.006 10.1016/j.mito.2017.10.006.

24. Martínez-Reyes, I., and Chandel, N.S. Mitochondrial TCA cycle metabolites control physiology and disease. 10.1038/s41467-019-13668-3.

25. Chevrollier, A., Loiseau, D., Reynier, P., and Stepien, G. (2011). Adenine nucleotide translocase 2 is a key mitochondrial protein in cancer metabolism. Biochimica et Biophysica Acta (BBA) -Bioenergetics 1807, 562–567. 10.1016/j.bbabio.2010.10.008.

26. Stevens, J.F., Revel, J.S., and Maier, C.S. (2018). Mitochondria-Centric Review of Polyphenol Bioactivity in Cancer Models. Antioxid Redox Signal 29, 1589–1611. 10.1089/ars.2017.7404.

27. Pham, A.H., Mccaffery, J.M., and Chan, D.C. (2012). Mouse lines with photo-activatable mitochondria to study mitochondrial dynamics. Genesis 50, 833–843. 10.1002/dvg.22050.

28. Zhang, D.J., Wang, Q., Wei, J., Baimukanova, G., Buchholz, F., Stewart, A.F., Mao, X., and Killeen, N. (2005). Selective Expression of the Cre Recombinase in Late-Stage Thymocytes Using the Distal Promoter of the *Lck* Gene. The Journal of Immunology 174, 6725–6731. 10.4049/jimmunol.174.11.6725.

29. van der Windt, G.J.W., Chang, C.H., and Pearce, E.L. (2016). Measuring bioenergetics in T cells using a seahorse extracellular flux analyzer. Curr Protoc Immunol 2016, 3.16B.1–3.16B.14. 10.1002/0471142735.im0316bs113.

30. van der Windt, G.J.W., Everts, B., Chang, C.H., Curtis, J.D., Freitas, T.C., Amiel, E., Pearce, E.J., and Pearce, E.L. (2012). Mitochondrial Respiratory Capacity Is a Critical Regulator of CD8 + T Cell Memory Development. Immunity 36, 68–78. 10.1016/j.immuni.2011.12.007.

31. Ahmad, T., Aggarwal, K., Pattnaik, B., Mukherjee, S., Sethi, T., Tiwari, B.K., Kumar, M., Micheal, A., Mabalirajan, U., Ghosh, B., et al. (2013). Computational classification of mitochondrial shapes reflects stress and redox state. Cell Death Dis 4, e461–e461. 10.1038/cddis.2012.213.

32. Hung, C.H.-L., Cheng, S.S.-Y., Cheung, Y.-T., Wuwongse, S., Zhang, N.Q., Ho, Y.-S., Lee, S.M.-Y., and Chang, R.C.-C. (2018). A reciprocal relationship between reactive oxygen species and mitochondrial dynamics in neurodegeneration. Redox Biol 14, 7–19. 10.1016/j.redox.2017.08.010.

33. Calvo, S.E., Clauser, K.R., and Mootha, V.K. (2016). MitoCarta2.0: An updated inventory of mammalian mitochondrial proteins. Nucleic Acids Res 44, D1251–D1257. 10.1093/nar/gkv1003.

34. Picca, A., and Lezza, A.M.S. (2015). Regulation of mitochondrial biogenesis through TFAM-mitochondrial DNA interactions. Useful insights from aging and calorie restriction studies. Preprint, 10.1016/j.mito.2015.10.001 10.1016/j.mito.2015.10.001.

35. Lin, S., Huang, C., Sun, J., Bollt, O., Wang, X., Martine, E., Kang, J., Taylor, M.D., Fang, B., Singh, P.K., et al. (2019). The mitochondrial deoxyguanosine kinase is required for cancer cell stemness in lung adenocarcinoma. EMBO Mol Med 11. 10.15252/emmm.201910849.

36. Inigo, M., Deja, S., and Burgess, S.C. (2021). Annual Review of Nutrition Ins and Outs of the TCA Cycle: The Central Role of Anaplerosis. 10.1146/annurev-nutr-120420.

37. Mak, T.W., Grusdat, M., Duncan, G.S., Dostert, C., Nonnenmacher, Y., Cox, M., Binsfeld, C., Hao, Z., Brüstle, A., Itsumi, M., et al. (2017). Glutathione Primes T Cell Metabolism for Inflammation. Immunity 46, 675–689. 10.1016/j.immuni.2017.03.019.

38. Harris, I.S., Treloar, A.E., Inoue, S., Sasaki, M., Gorrini, C., Lee, K.C., Yung, K.Y., Brenner, D., Knobbe-Thomsen, C.B., Cox, M.A., et al. (2015). Glutathione and Thioredoxin Antioxidant Pathways Synergize to Drive Cancer Initiation and Progression. Cancer Cell 27, 211–222. 10.1016/j.ccell.2014.11.019.

39. Ju, H.Q., Lin, J.F., Tian, T., Xie, D., and Xu, R.H. (2020). NADPH homeostasis in cancer: functions, mechanisms and therapeutic implications. Preprint at Springer Nature, 10.1038/s41392-020-00326-0 10.1038/s41392-020-00326-0.

40. Ma, E.H., Bantug, G., Griss, T., Condotta, S., Johnson, R.M., Samborska, B., Mainolfi, N., Suri, V., Guak, H., Balmer, M.L., et al. (2017). Serine Is an Essential Metabolite for Effector T Cell Expansion. Cell Metab 25, 345–357. 10.1016/j.cmet.2016.12.011.

41. Ron-Harel, N., Notarangelo, G., Ghergurovich, J.M., Paulo, J.A., Sage, P.T., Santos, D., Kyle Satterstrom, F., Gygi, S.P., Rabinowitz, J.D., Sharpe, A.H., et al. (2018). Defective respiration and one-carbon metabolism contribute to impaired naïve T cell activation in aged mice. Proc Natl Acad Sci U S A 115, 13347–13352. 10.1073/pnas.1804149115.

42. Dutt, S., Hamza, I., and Bartnikas, T.B. (2022). Molecular Mechanisms of Iron and Heme Metabolism. Annu Rev Nutr 42, 311–335. 10.1146/annurev-nutr-062320-112625.

43. Berg, V., Modak, M., Brell, J., Puck, A., Künig, S., Jutz, S., Steinberger, P., Zlabinger, G.J., and Stöckl, J. (2020). Iron Deprivation in Human T Cells Induces Nonproliferating Accessory Helper Cells. Immunohorizons 4, 165–177. 10.4049/immunohorizons.2000003.

44. Jennifer, B., Berg, V., Modak, M., Puck, A., Seyerl-Jiresch, M., Künig, S., Zlabinger, G.J., Steinberger, P., Chou, J., Geha, R.S., et al. (2020). Transferrin receptor 1 is a cellular receptor for human heme-albumin. Commun Biol 3, 621. 10.1038/s42003-020-01294-5.

45. Teh, M.R., Frost, J.N., Armitage, A.E., and Drakesmith, H. (2021). Analysis of Iron and Iron-Interacting Protein Dynamics During T-Cell Activation. Front Immunol 12. 10.3389/fimmu.2021.714613.

46. Zhou, Y., Zhou, B., Pache, L., Chang, M., Khodabakhshi, A.H., Tanaseichuk, O., Benner, C., and Chanda, S.K. (2019). Metascape provides a biologist-oriented resource for the analysis of systems-level datasets. Nat Commun 10. 10.1038/s41467-019-09234-6.

47. Samborska, B., Roy, D.G., Rahbani, J.F., Hussain, M.F., Ma, E.H., Jones, R.G., and Kazak, L. (2022). Creatine transport and creatine kinase activity is required for CD8+ T cell immunity. Cell Rep 38. 10.1016/j.celrep.2022.110446.

48. Di Biase, S., Ma, X., Wang, X., Yu, J., Wang, Y.C., Smith, D.J., Zhou, Y., Li, Z., Kim, Y.J., Clarke, N., et al. (2019). Creatine uptake regulates CD8 T cell antitumor immunity. Journal of Experimental Medicine 216, 2869–2882. 10.1084/JEM.20182044.

49. Greenhaff, P.L. (2001). The creatine-phosphocreatine system: there’s more than one song in its repertoire. J Physiol 537, 657–657. 10.1111/j.1469-7793.2001.00657.x.

50. Esterhuizen, K., van der Westhuizen, F.H., and Louw, R. (2017). Metabolomics of mitochondrial disease. Mitochondrion 35, 97–110. 10.1016/J.MITO.2017.05.012.

51. Westbrook, R.L., Bridges, E., Roberts, J., Escribano-Gonzalez, C., Eales, K.L., Vettore, L.A., Walker, P.D., Vera-Siguenza, E., Rana, H., Cuozzo, F., et al. (2022). Proline synthesis through PYCR1 is required to support cancer cell proliferation and survival in oxygen-limiting conditions. Cell Rep 38, 110320. 10.1016/J.CELREP.2022.110320.

52. Schwörer, S., Berisa, M., Violante, S., Qin, W., Zhu, J., Hendrickson, R.C., Cross, J.R., and Thompson, C.B. (2020). Proline biosynthesis is a vent for TGFβ-induced mitochondrial redox stress. EMBO J 39. 10.15252/EMBJ.2019103334.

53. Phang, J.M. (2012). The Proline Regulatory Axis and Cancer. Front Oncol 0, 60. 10.3389/FONC.2012.00060.

54. Phang, J.M. Proline Metabolism in Cell Regulation and Cancer Biology: Recent Advances and Hypotheses. 10.1089/ars.2017.7350.

55. Yang, Z., Zhao, X., Shang, W., Liu, Y., Ji, J.-F., Liu, J.-P., and Tong, C. (2020). Pyrroline-5-carboxylate synthase senses cellular stress and modulates metabolism by regulating mitochondrial respiration. Cell Death & Differentiation 2020 28:1 28, 303–319. 10.1038/s41418-020-0601-5.

56. Phang, J.M., Downing, S.J., Yeh, G.C., Smith, R.J., Williams, J.A., and Hagedorn, C.H. (1982). Stimulation of the hexosemonophosphate-pentose pathway by pyrroline-5-carboxylate in cultured cells. J Cell Physiol 110, 255–261. 10.1002/jcp.1041100306.

57. Balmer, M.L., and Hess, C. (2017). Starving for survival—how catabolic metabolism fuels immune function. Preprint at Elsevier Ltd, 10.1016/j.coi.2017.03.009 10.1016/j.coi.2017.03.009.

58. Owen, O.E., Kalhan, S.C., and Hanson, R.W. (2002). The Key Role of Anaplerosis and Cataplerosis for Citric Acid Cycle Function *. Journal of Biological Chemistry 277, 30409–30412. 10.1074/JBC.R200006200.

59. Varanasi, S.K., Ma, S., and Kaech, S.M. (2019). T Cell Metabolism in a State of Flux. Preprint at Cell Press, 10.1016/j.immuni.2019.10.012 10.1016/j.immuni.2019.10.012.

60. Elia, I., Rowe, J.H., Johnson, S., Joshi, S., Notarangelo, G., Kurmi, K., Weiss, S., Freeman, G.J., Sharpe, A.H., and Haigis, M.C. (2022). Tumor cells dictate anti-tumor immune responses by altering pyruvate utilization and succinate signaling in CD8+ T cells. Cell Metab 34, 1137–1150.e6. 10.1016/J.CMET.2022.06.008.

61. Kiesel, V.A., Sheeley, M.P., Coleman, M.F., Cotul, E.K., Donkin, S.S., Hursting, S.D., Wendt, M.K., and Teegarden, D. (2021). Pyruvate carboxylase and cancer progression. Cancer Metab 9. 10.1186/S40170-021-00256-7.

62. Menk, A. V., Scharping, N.E., Moreci, R.S., Zeng, X., Guy, C., Salvatore, S., Bae, H., Xie, J., Young, H.A., Wendell, S.G., et al. (2018). Early TCR Signaling Induces Rapid Aerobic Glycolysis Enabling Distinct Acute T Cell Effector Functions. Cell Rep 22, 1509–1521. 10.1016/j.celrep.2018.01.040.

63. Luciani, S., Martini, N., and Santi, R. (1971). Effects of carboxyatractyloside a structural analogue of atractyloside on mitochondrial oxidative phosphorylation. Life Sci 10, 961–968. 10.1016/0024-3205(71)90099-3.

64. Zhang, P., Cheng, X., Sun, H., Li, Y., Mei, W., and Zeng, C. (2021). Atractyloside Protect Mice Against Liver Steatosis by Activation of Autophagy via ANT-AMPK-mTORC1 Signaling Pathway. Front Pharmacol 12. 10.3389/fphar.2021.736655.

65. Cho, J., Zhang, Y., Park, S.-Y., Joseph, A.-M., Han, C., Park, H.-J., Kalavalapalli, S., Chun, S.-K., Morgan, D., Kim, J.-S., et al. (2017). Mitochondrial ATP transporter depletion protects mice against liver steatosis and insulin resistance. Nat Commun 8, 14477. 10.1038/ncomms14477.

66. Permyakova, A., Hamad, S., Hinden, L., Baraghithy, S., Kogot-Levin, A., Yosef, O., Shalev, O., Tripathi, M.K., Amal, H., Basu, A., et al. (2024). Renal Mitochondrial ATP Transporter Ablation Ameliorates Obesity-Induced CKD. Journal of the American Society of Nephrology 35, 281–298. 10.1681/ASN.0000000000000294.

67. MacIver, N.J., Michalek, R.D., and Rathmell, J.C. (2013). Metabolic Regulation of T Lymphocytes. Annu Rev Immunol 31, 259–283. 10.1146/annurev-immunol-032712-095956.

68. Broeks, M.H., Meijer, N.W.F., Westland, D., Zwartkruis, F.J.T., Verhoeven-Duif, N.M., Jans, J.J.M., Bosma, M., Gerrits, J., German, H.M., Ciapaite, J., et al. (2023). The malate-aspartate shuttle is important for de novo serine biosynthesis ll The malate-aspartate shuttle is important for de novo serine biosynthesis. Cell Rep 42. 10.1016/j.celrep.2023.113043.

69. Seo, J.B., Riopel, M., Cabrales, P., Huh, J.Y., Bandyopadhyay, G.K., Andreyev, A.Y., Murphy, A.N., Beeman, S.C., Smith, G.I., Klein, S., et al. (2019). Knockdown of ANT2 reduces adipocyte hypoxia and improves insulin resistance in obesity. Nat Metab 1, 86–97. 10.1038/s42255-018-0003-x.

70. Moon, J.S., da Cunha, F.F., Huh, J.Y., Andreyev, A.Y., Lee, J., Mahata, S.K., Reis, F.C.G., Nasamran, C.A., and Lee, Y.S. (2021). ANT2 drives proinflammatory macrophage activation in obesity. JCI Insight 6. 10.1172/jci.insight.147033.

71. Kokoszka, J.E., Waymire, K.G., Flierl, A., Sweeney, K.M., Angelin, A., MacGregor, G.R., and Wallace, D.C. (2016). Deficiency in the mouse mitochondrial adenine nucleotide translocator isoform 2 gene is associated with cardiac noncompaction. Biochimica et Biophysica Acta (BBA) -Bioenergetics 1857, 1203– 1212. 10.1016/j.bbabio.2016.03.026.

72. Murphy, M.P., and Siegel, R.M. (2013). Mitochondrial ROS Fire Up T Cell Activation. Preprint, 10.1016/j.immuni.2013.02.005 10.1016/j.immuni.2013.02.005.

73. Sena, L.A., Li, S., Jairaman, A., Prakriya, M., Ezponda, T., Hildeman, D.A., Wang, C.R., Schumacker, P.T., Licht, J.D., Perlman, H., et al. (2013). Mitochondria Are Required for Antigen-Specific T Cell Activation through Reactive Oxygen Species Signaling. Immunity 38, 225–236. 10.1016/j.immuni.2012.10.020.

74. Chapman, N.M., Boothby, M.R., and Chi, H. (2020). Metabolic coordination of T cell quiescence and activation. Preprint at Nature Research, 10.1038/s41577-019-0203-y 10.1038/s41577-019-0203-y.

75. Dimeloe, S., Burgener, A.V., Grählert, J., and Hess, C. (2017). T-cell metabolism governing activation, proliferation and differentiation; a modular view. Preprint at Blackwell Publishing Ltd, 10.1111/imm.12655 10.1111/imm.12655.

